# Entrenchment and contingency in neutral protein evolution with epistasis

**DOI:** 10.1101/2025.01.09.632266

**Authors:** Lisa Schmelkin, Vincenzo Carnevale, Allan Haldane, Jeffrey P. Townsend, Sarah Chung, Ronald M. Levy, Sudhir Kumar

## Abstract

Protein sequence evolution in the presence of epistasis makes many previously acceptable amino acid residues at a site unfavorable over time. This phenomenon of entrenchment has also been observed with neutral substitutions using Potts Hamiltonian models. Here, we show that simulations using these models often evolve non-neutral proteins. We introduce a Neutral-with-Epistasis (N×E) model that incorporates purifying selection to conserve fitness, a requirement of neutral evolution. N×E protein evolution revealed a surprising lack of entrenchment, with site-specific amino-acid preferences remaining remarkably conserved, in biologically realistic time frames despite extensive residue coupling. Moreover, we found that the overdispersion of the molecular clock is caused by rate variation across sites introduced by epistasis in individual lineages, rather than by historical contingency. Therefore, substitutional entrenchment and rate contingency may indicate that adaptive and other non-neutral evolutionary processes were at play during protein evolution.

## Introduction

The Neutral Theory of Molecular Evolution (NTME) posits that most observed protein sequence differences are due to the random fixation of selectively neutral alleles in orthologous proteins in different species (*1-3*). Neutral patterns of accumulation of these variations are commonly used as a baseline to detect molecular signatures of adaptive evolution and natural selection. However, theoretical analyses and tests of neutral evolution have been developed with a simplifying assumption of independence of evolutionary substitutions across sites in a sequence, despite general appreciation that interactions among amino acid residues (epistasis) preserve the structure and function of proteins.

Building protein evolutionary models incorporating epistasis has been challenging because correlations between sites arising from shared evolutionary history and limited variation among protein sequences cannot be disentangled from actual direct couplings between them with a few sequences. With the introduction of Direct Coupling Analysis (DCA), a sequence co-variation model based on a Potts Hamiltonian energy function, epistasis can be readily modeled and incorporated into molecular evolutionary studies using large alignments (*4*). Using DCA, it has been reported that epistasis causes large differences in evolutionary rates over time and entrenchment of amino acid substitutions during neutral evolution within ancient protein families (*5*), as well as more generally in non-neutral evolution or evolution within viral populations (*6-12*).

If neutral substitutions in the presence of epistasis cause significant substitutional entrenchment, then the presence of entrenchment cannot be used as a sign of beneficial and adaptive molecular evolutionary substitutions, which are notoriously difficult to identify. This prompted us to investigate the claims of substitution entrenchment and overdispersion of evolutionary rates during the neutral evolution of proteins. Specifically, a Sequence Evolution with Epistatic Contributions (hereafter, SEEC) framework was developed for protein evolutionary simulations using a Potts Hamiltonian (PH) model to study the emergent properties of neutral evolution of domain families (*5*). DCA was used to produce PH models of epistasis between amino acid residues across sites using multiple sequence alignments (MSAs) composed of thousands of homologous protein domains (*13*).

In a PH model, every pair of sites has an amino-acid residue interaction matrix, e.g., sites 121 and 133 in domains belonging to the family PF00001 and inferred using an MSA containing 26,346 raw sequences (**Fig. 1**). For example, in this matrix, 121Trp is preferred (blue) and 121Asp is avoided (red) when an Ala residue occupies position 133. The color intensity indicates the strength (magnitude) of the interaction, and there are many favorable or unfavorable site-residue combinations. The SEEC framework was employed to examine emergent patterns of neutral molecular evolution in the presence of epistasis (*5, 14*). SEEC simulations result in variability of amino acid substitution rates over time and among sites as well as evolutionary Stokes’ shift (*5, 14*).

**Figure 1.**
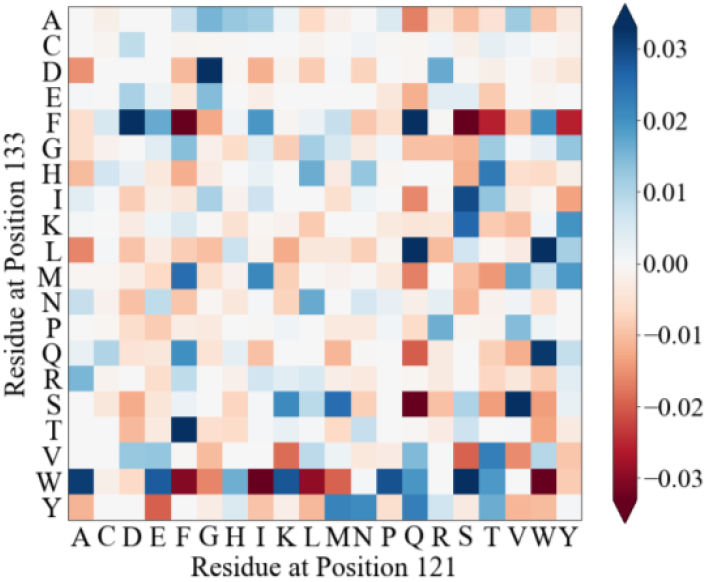
Pairwise couplings captured by the PH model. Amino-acid residue couplings for one pair of sites (121 and 133) in the Potts Hamiltonian model for the domain family PF00001, estimated by de la Paz et al. (*5*). Positive values (blue) indicate commonly observed combinations in the MSA of 26,346 PF0001 domain sequences, whereas negative values (red) correspond to unpreferred residue combinations. Individual values correspond to Direct Information (*4*).

However, a fundamental assumption in SEEC and similar PH simulation frameworks is that the amino-acid substitutions permitted are neutral. This key assumption is yet to be tested, which is needed because the PH models used in SEEC are inferred from large and diverse MSAs of homologs, including proteins with various functions. For example, the PH model fit to the PF00001 domain family was derived from 26,346 sequences found in G-protein-coupled receptor proteins (*5*). Proteins contributing homologous functional domains in this MSA serve vastly different roles, such as hormones, neurotransmitters, and light receptors. Many of these proteins have evolved by gene duplications, producing paralogous proteins and domains. Consequently, PH models inferred from such MSAs will incorporate many amino acid substitutions between paralogous proteins with vastly different functions. Therefore, these PH models may allow non-neutral amino acid substitutions, i.e., those not permissible when considering only orthologous sequences and neutral evolution. As a result, the reported patterns of evolutionary entrenchment and rate variability among lineages may be due to the introduction of non-neutral substitutions during protein evolution.

This article presents results from our investigation of the neutrality of protein evolution permitted by the SEEC framework. We compare properties of orthologs and non-orthologs under PH models fit using MSAs effectively composed of paralogs (see *Methods*). Upon finding non-neutral evolutionary patterns, we developed an approach to simulate neutral sequence evolution with substitutional epistasis, referred to as the Neutral-with-Epistasis (N×E) framework. Then, we used N×E to test if neutral evolution can cause patterns of evolutionary rate variability and substitutional entrenchment.

## Results and Discussion

### Assessing the neutrality of SEEC evolution

From the collection of domain families analyzed previously using SEEC (*5*), we selected the PF00001 domain family because it contains the longest sequences (up to 268 residues). The MSA for PF00001 contains 26,346 sequences that were previously used to infer the PH model (*Methods*). To test the orthology of sequences and evaluate the neutrality of amino acid substitutions permitted by SEEC, we retrieved a collection of amino-acid sequence alignments of vertebrate orthologs from the UCSC resource (*15*). We extracted the intersection of human proteins in the UCSC MSAs with that of the PF00001 domain sequences. This analysis yielded 80 proteins for which a sequence in the PF00001 MSA matched perfectly with a segment of the human sequence in the UCSC alignments (see *Methods*). Portions of the multi-species UCSC alignments corresponding to the matched domain were retained, producing 80 ortho-domain alignments, each with sequences of up to 100 vertebrate species ranging from humans to fish. **Figure 2** shows an alignment for the 5-hydroxytryptamine receptor 5A (HTR5A) ortho-domain. Pairwise comparison of sequences within this alignment showed less than 10% sequence difference, on average (mean *p*-distance < 0.1; **Fig. 3a**).

**Figure 2.**
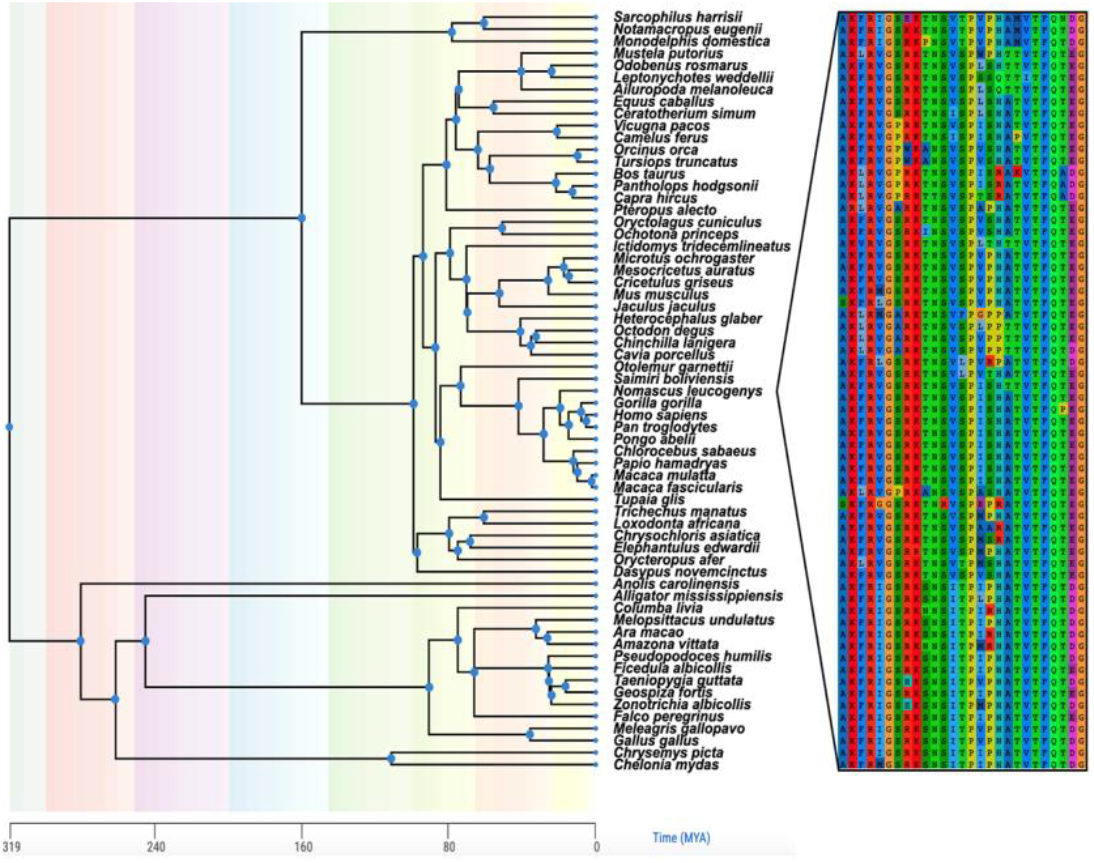
Phylogeny of vertebrate species in the HTR5A ortho-domain alignment. An alignment of PF00001 domains found in the HTR5A protein was constructed. Sixty-four species remained after removing sequences with indels or missing data. Species divergence times were retrieved from Timetree.org. A snippet of positions demonstrating sequence variation among species is shown beside the phylogeny

**Figure 3.**
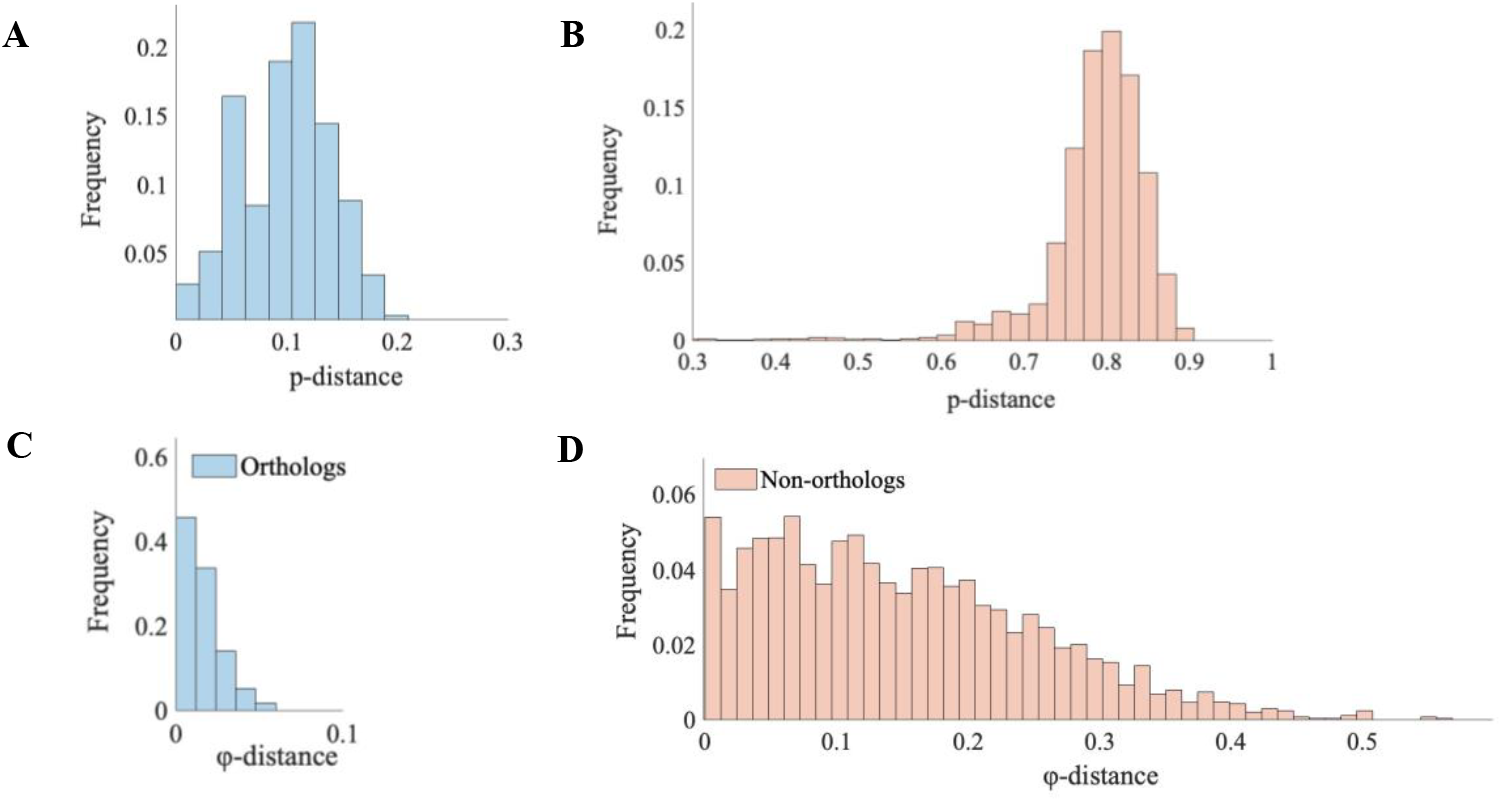
(**A**) *p*-distance among pairs of orthologous sequences (blue; domains found in the HTR5A protein) compared to (**B**) *p*-distance among pairs of non-orthologs (red; inter-domain comparisons). Note that *p*-distance is equivalent to the Hamming distance which compares strings of equal length. The dataset of non-orthologs was constructed by randomly selecting one sequence from each of the 80 ortho-domain alignments (see *Methods*) (**C**) φ-distance among PF00001 orthologs found in the HTR5A protein. For every pair of species in the ortho-domain, φ-distance was calculated as a change in energy (Δφ) divided by the mean φ of the sequence pair. Bars in the histogram show the frequency of φ-distances among all possible species pairings. (**D**) φ-distance among pairs of non-orthologs (red; inter-domain comparisons).

This pattern held for other ortho-domains (**Fig. S1a**); 90% of the 80 ortho-domain alignments had a maximum *p*-distance of less than 0.3 (**Fig. S2**), which we consider a biologically realistic maximum limit of domain divergence in vertebrates.

In contrast, sequence comparisons across ortho-domains (non-orthologs) yielded an average *p*-distance of 0.8 (**Fig. 3b**). The smallest *p*-distance between sequences from different ortho-domains was 0.3, which was much larger than any *p*-distance between sequences from the same ortho-domain. Therefore, PF00001 proteins that have evolved due to speciation and, thus, expected to retain the same function, have domain homologs with more similar sequences than non-orthologous domains. Many of the latter domains in the PF00001 family likely evolved by gene duplications or harbor only structural similarity, which likely experienced non-neutral substitutions.

Next, we evaluated the fitness of sequences found across species within each ortho-domain alignment. We used Potts Hamiltonian Energy (PHE; φ) as a proxy for the biophysical properties, function, and fitness of individual domain sequences. This is because φ is well-correlated with the free energy of protein folding as well as many empirical measurements, such as gene fitness, antibiotic resistance, and viral replicative capacity (*13, 16-21*).

We tested the primary assumption of the neutral theory that sequences within ortho-domains will have a similar φ because they reside in orthologous proteins with a similar function among species. We calculated φ for all the sequences in 80 ortho-domain alignments. Then, we normalized φ-distances for pairs of sequences within and between ortho-domain alignments, where φ-distance is the difference in φ between domains divided by the average φ of the domains compared. The distribution of φ-distances within the HTR5A ortho-domain alignment (orthologs) is narrow with a mean of 0.016 (**Fig. 3c**). Low φ-distances were also observed within other ortho-domains (**Fig. S1b**: distribution of average φ-distance within ortho-domains). In contrast, the average φ-distance between non-orthologs is about ten times larger (0.150; **Fig. 3d**). Therefore, φ is well conserved within ortho-domains but not among non-orthologs, which is consistent with the neutral theory of molecular evolution.

We found that the range of φ varied across ortho-domains. While some ortho-domains had more variation in energy among species than others, the ranges of φ were typically narrow: within ± 2.4% of the mean on average (**Fig. 4**). While it would be simplistic to assume that there is a one-to-one relationship between φ and biological fitness, these results clearly show that orthologous domain sequences are energetically similar. This interpretation is consistent with the empirical evidence that shows φ is conserved in proteins with the same function. For example, in the analysis of lysin catalytic domain sub-libraries, φ was similar among proteins with the same function. In fact, the fraction of stable variants quickly dropped to 0% when φ was not conserved (*22*). Such a drop-off was also seen for in vitro fitness measurements of HIV-1 strains containing Gag mutations, where small deviations in φ resulted in zero replicative capacity of the virus (*23*).

**Figure 4.**
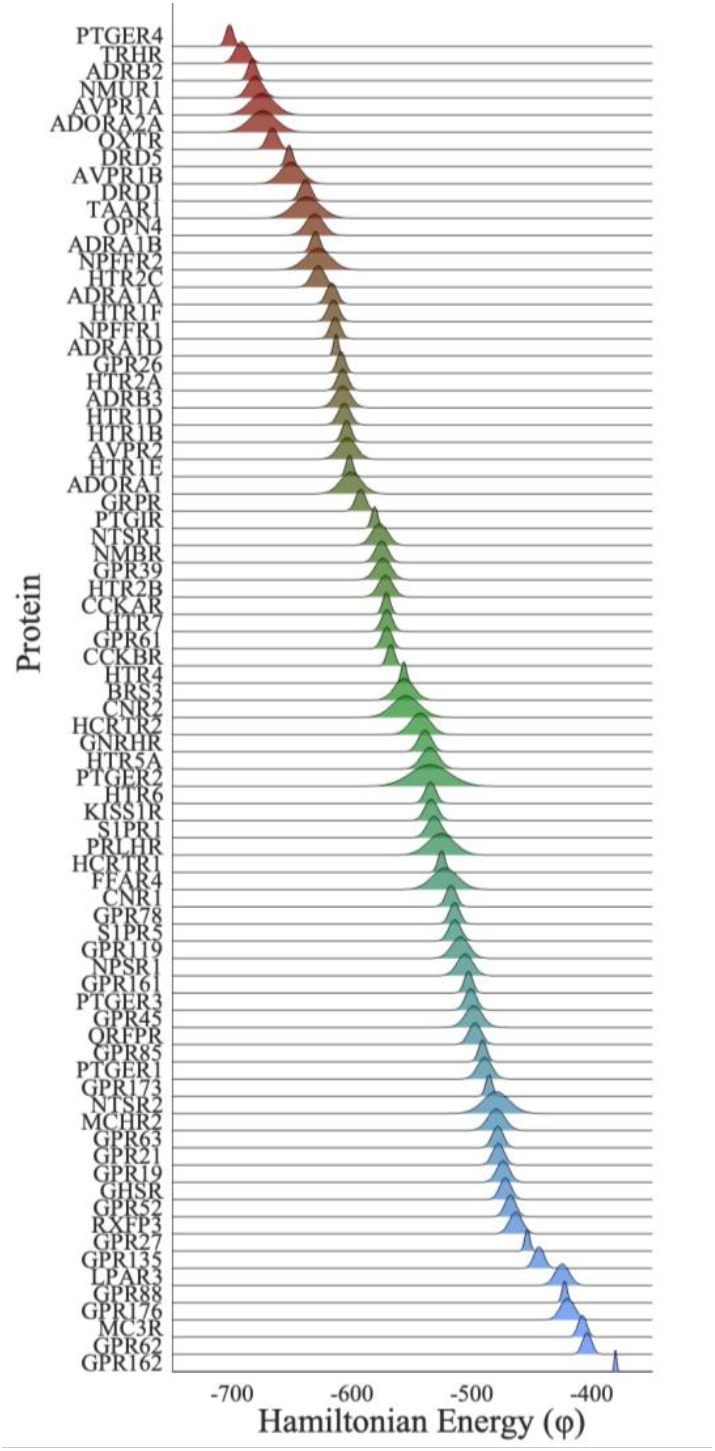
Summary of φ values within 78 ortho-domain alignments of PF00001. The y-axis indicates the protein in which the domain is found (HGNC official symbols are used). For each ortho-domain, a normalized distribution of energies (φ) is shown,3w5 here the domains are sorted by mean energy.

### Domain φ trajectories during protein evolution

The large difference in φ among domains included in the PF00001 MSA has consequences for the neutrality of evolution permitted by SEEC. **Fig. 5** shows the trajectories of φ for domains evolved by the SEEC model, where the energy wanders in and out of the neutral zone as substitutions accumulate. This figure shows five simulation replicates, each starting with the same sequence of a PF00001 domain found in melanocortin 3 receptor (MC3R; |φ| = 408). Each domain was evolved to an evolutionary distance *d* = 0.25 substitutions per site, which resulted in a 30% sequence difference between the initial and the final sequences (average *p*-distance = 0.30). This average is similar to the maximum *p*-distance observed within ortho-domain alignments (**Fig. S2**). Therefore, the evolution simulated here mirrors protein sequence evolution observed among diverse vertebrates. Across all simulated trajectories, SEEC evolved a non-neutral domain, i.e., one with φ outside the empirically observed range of empirical MC3R domains (−415 to -400), taking as few as 3 substitutions.

**Figure 5.**
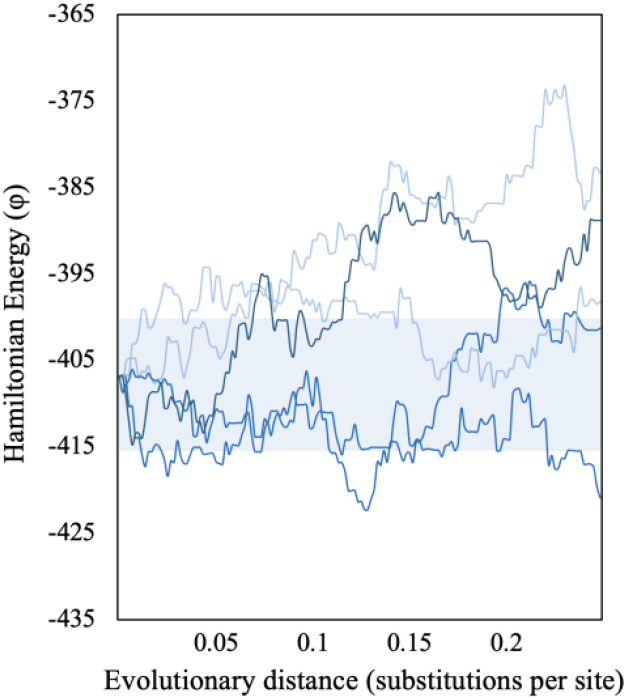
Trajectories of φ for SEEC lineages evolved to an evolutionary distance (d) of 0.25 substitutions per site. All 5 simulated lineages begin with the same domain sequence found in the 2M0C3R protein. The shaded region indicates the neutral zone (φ= -415 to -400).

The proportion of evolved sequences in the neutral zone for PF000001 domains with low (380.7 < |φ| < 514.5; 26 domains), median (514.7 < |φ| < 601.0; 26 domains), and high PH energies (602.2 < |φ| < 702.1; 26 domains) decreased with time. Even with limited sequence divergence, i.e., early in evolutionary history, many evolved domains are no longer neutral based on their PH energies. As more substitutions accumulate, the evolving domain tends to escape the neutral zone more frequently, regardless of the energy of the starting sequence (**Fig. 6**).

**Figure 6.**
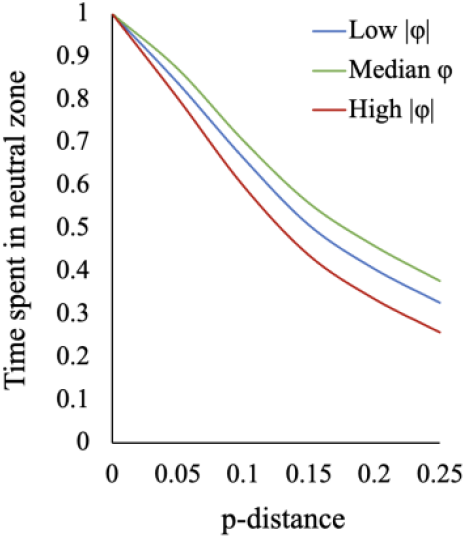
Time spent in the neutral zone for SEEC simulations. (**A**) Simulations beginning from ortho-domain sequences with low energy (−380.7 to -514.5), median energy (−514.7 to -601.0), and high energy (−602.2 to -702.1). At increasing *p*-distances of 0.05 - 0.25, the proportion of time spent in the neutral zone was recorded. 100 lineages were simulated for each ortho-domain.

A simple explanation for the non-neutrality of SEEC evolution observed here is that the same PH model is applied to all the domains, i.e., epistatic patterns are homogenous among domains. This is necessitated to do statistical inference of pairwise couplings. PH models utilized here were inferred using bmDCA and are thought to best capture many aspects of epistasis (see *Methods*; *24*). However, as noted above, the domain collection used to infer the PH model is from proteins with highly diverse functions, which may have evolved protein- and function-specific epistatic patterns. It is clearly the case that different ortho-domains, which frequently show no overlap in PH energy with other ortho-domains, have specific patterns in epistasis to preserve the function of the proteins in which they are found (**Fig. 4**). Therefore, a single PH model will not fit the evolution of some (or any) of the ortho-domains, which means that preferred amino-acid substitutions based on a common PH model can indeed be non-neutral.

### A Neutral-with-Epistasis (NxE) model of domain evolution

The non-neutral evolution observed for PH models inferred from domains with sequence and structural homology indicates that the use of PH models inferred from alignments of orthologous sequences would be ideal for simulating neutral evolution using SEEC. This inference is not feasible from species-level sequence data because the number of sequences in each ortho-alignment and substitutions therein is far too few to distinguish direct pairwise couplings between site-residue pairs and correlation caused by common ancestry. To overcome this limitation, we took an alternate approach in which the global PH model was used in SEEC, but the descendant domains failing to conserve φ were further subjected to purifying selection. That is, a substitution permitted by the PH model in SEEC was considered neutral if and only if it resulted in a sequence with φ within the range observed in the ortho-domain alignment from which the ancestral sequence was drawn. This Neutral-with-Epistasis (N×E) framework was implemented by modifying the SEEC source code. Essentially, N×E only allows amino acid substitutions that are permissible by the PH model and fall within the neutral φ range (fitness conservation). **Fig. 7** shows the flow of data analysis and simulations performed using N×E. In our simulations, the empirically derived limits on φ for amino acid substitutions were those presented in **Fig. 4** (see also **Table S1** and *Methods*).

**Figure 7.**
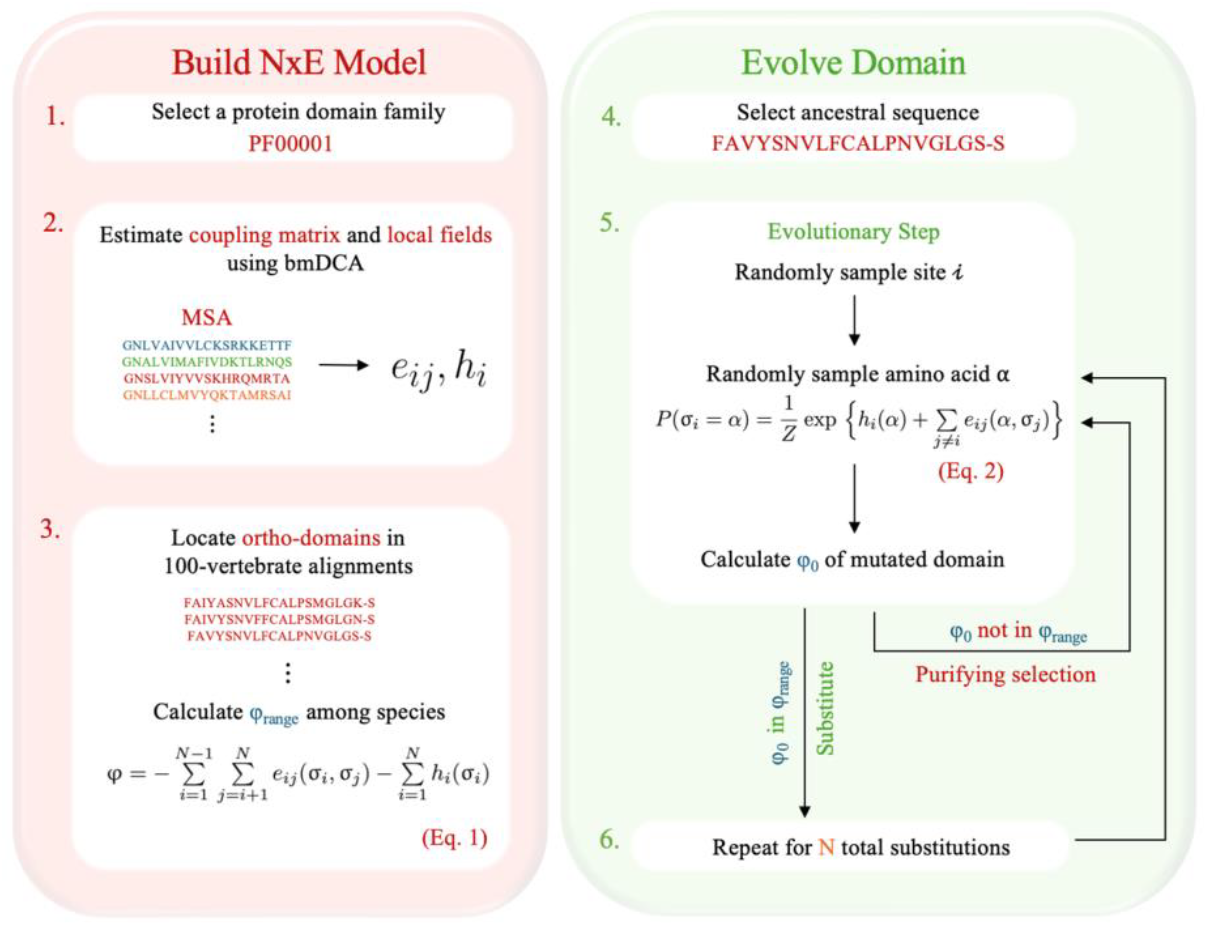
The N×E schematic. (1–2) We first select a protein domain family and estimate PH model parameters using bmDCA: a coupling matrix e for each pair of sites, and local fields h, or site-specific amino-acid preferences (*25*) (3) We then run a BLAST search of all domain sequences against a database of human protein-coding genes selected from the UCSC 100 vertebrate alignments (*15*), and calculate a range in φ for orthologs of each domain. (4) We select a sequence native to the ortho-domain and simulate neutral sequence evolution. (5) At each evolutionary step, we randomly select a position along the protein from a uniform distribution and then sample a new residue from a conditional probability distribution, inferred from the statistics of the MSA of the family (see *Methods*). We calculate the new φ and impose purifying selection against all substitutions that are predicted to be non-neutral. We instead sample a new site and residue as necessary until we find a permissible substitution (or retain the same residue). (6) We repeat this procedure for a specified number of evolutionary steps or the total number of substitutions.

### Emergent properties of molecular evolution under N×E

#### Contingency of evolutionary rates

Excess temporal variation in evolutionary rates has been reported for sequences simulated using PH models in single lineages (*5*). Instead, we used a 100-species star phylogeny, a classical setup to examine rate variation among lineages (see *Methods* for details) (*3, 26, 27*). An overdispersion of evolutionary rates (*R* > *v*/*m*) among lineages was found in N×E simulations as well, i.e., the variance (*v*) of lineage rates exceeded the mean (*m*). But *R* in N×E is relatively modest and plateaus quickly (**Fig. 8a**) as opposed to the large *R* reported previously with SEEC (*5*). Such trends are also observed for other PF00001 ortho-domains (**Table S2**).

**Figure 8.**
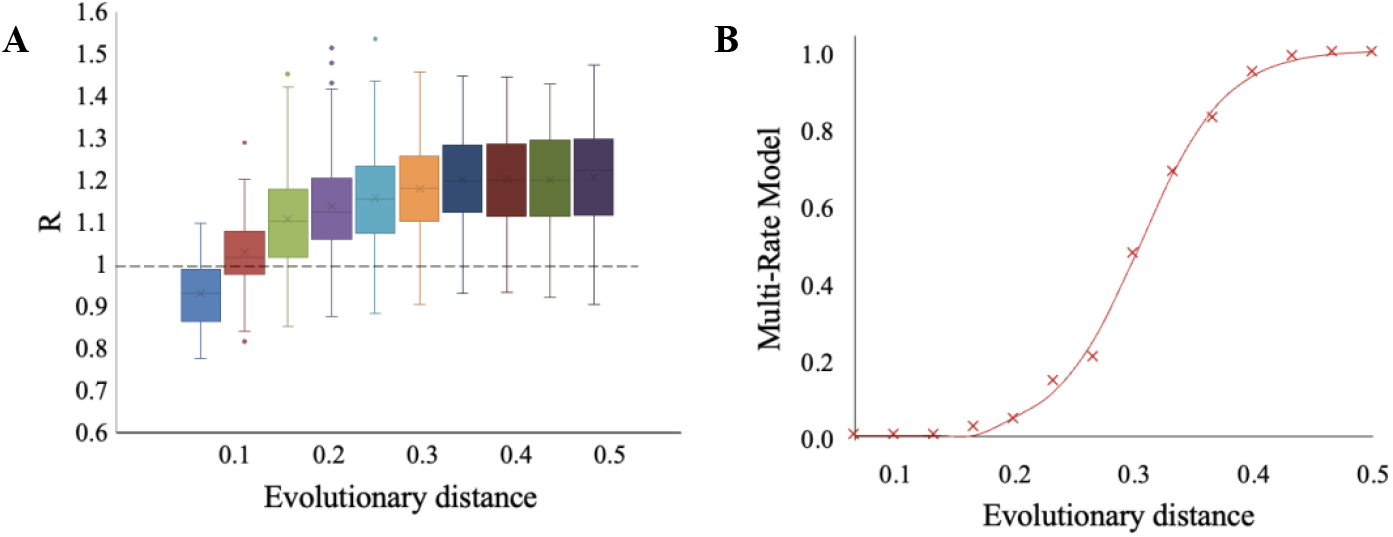
Overdispersion (*R* > 1) of evolutionary rates in N×E lineages. (**A**) Box-and-whisker plots of *R*, the ratio of the variance to the mean of substitution counts in 100-tip star phylogenies. One hundred replicates were simulated using the PF00001 ortho-domain found in the PTGER4 protein until evolutionary distances of *d* = 0.1, 0.2, 0.3, 0.4, and 0.5 substitutions per site, on average. The dotted line indicates *R* = 1 which is the expectation when substitution rates are uniform across lineages. (**B**) Rate-among-sites model fit of 100 N×E lineages (PF00001 domain, PTGER4 protein). Four models were considered: single-rate (S), gamma function (Γ), single-rate with invariant sites (I+S), or a gamma function with invariant sites (I+Γ). The best model was determined by Akaike information criterion (AICc) values (*14, 28, 29*). The y axis shows the proportion of lineages in which a multi-rate model (I+S, Γ, I+Γ) provided the best fit, for lineages evolved to the same evolutionary distances as panel A.

One explanation for persistent rate variation (*R* > 1) is the contingency of evolutionary rates, such that a slower-evolving lineage will tend to evolve slowly in the future, whereas a faster-evolving lineage will continue evolving faster. In other words, some neutral substitutions may speed up evolution early in a lineage, predisposing that lineage to continue to evolve quickly for at least some time. We tested contingency in evolutionary rates using a runs-test for every lineage in the star tree. Evolutionary rates were sampled over 10 epochs with the same number of evolutionary steps, with an average of 0.05 substitutions per site in each epoch. If there are multiple consecutive epochs with either fast or slow evolutionary rates in the same direction, this would suggest contingency with *P* < 0.05 used to reject the null hypothesis of no contingency. At a 5% significance level, fewer than 3% of all lineages rejected the null hypothesis in an analysis of all variable PF00001 ortho-domains (**Fig. 9**). Therefore, results are consistent with the null expectation of no contingency.

**Figure 9.**
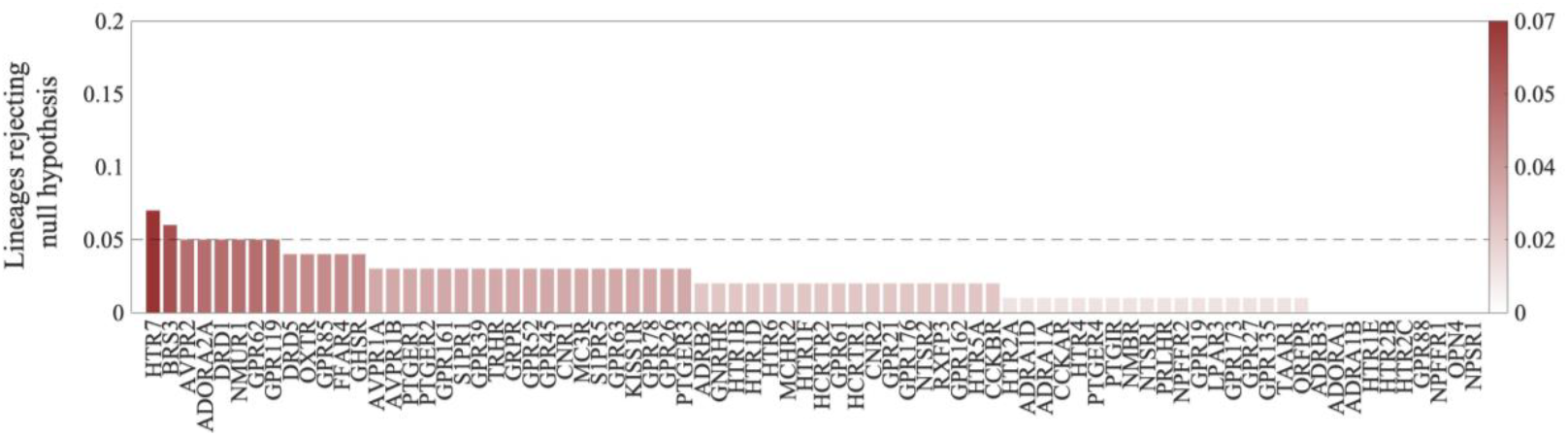
Contingency of evolutionary rates in N×E evolution. Percentage of lineages in a star tree showing contingency for all 78 variable ortho-domains. For each domain, a star tree was simulated consisting of 100 lineages, and the relative evolutionary rate was found for each lineage in epochs of 0.05 substitutions per site. A runs test was performed on each lineage to assess whether relative evolutionary rates in each epoch were mutually independent (random). The percentage of lineages with *P* < 0.05 is shown for each ortho-domain.

An examination of replicates with a *P* < 0.05 revealed that the rate oscillation between slow and fast evolving from one epoch to the next was the primary reason for rejecting the null hypothesis rather than contingency (**Fig. S4**). Therefore, we did not detect any significant runs of decreased or increased rates over time, meaning that past evolutionary rates do not determine the rate trajectory of future changes. That is, neutral evolution with epistasis does not result in the contingency of evolutionary rates.

Instead, the observed overdispersion of evolutionary rates is explained by the variation in rates caused by epistasis among positions of a domain (e.g., (*14*)). We, therefore, tested the hypothesis that a multi-rate model could explain the modest overdispersion (*R* > 1) observed in N×E simulations. We compared the fit of a single-rate (S) model with a set of multi-rate (MR) models by using Akaike information criterion (AICc) values (*28, 29*) as applied in a previous study (*14*) (see **Fig. 8** legend). In a star tree consisting of 100 lineages, we scored the fraction of lineages in which the MR model fit the site frequency distribution of the number of substitutions (*f*), which is shown in **Fig. 8b**. While *f* increases slowly in the early phases of molecular evolution, since only a few substitutions are permitted, it begins to grow significantly around *d* = 0.2 and reaches 100% concurrently with the plateauing of *R* (compare **Fig. 8a** and **8b**). This suggests that the introduction of significant rate variation among positions in individual lineages, due to greater variance than mean for the number of substitutions in individual lineages, causes an overdispersion of molecular clocks.

#### Entrenchment of neutral substitutions

Epistasis is routinely reported to cause entrenchment in protein evolution (*5, 15, 30*). That is, an amino acid residue that was previously acceptable at a particular site begins to be avoided after substitutions at other positions. Consequently, entrenchment can alter site-specific amino-acid preferences in descendants as compared to the ancestral protein due to epistasis. It is a common procedure to detect entrenchment by estimating the energy (fitness) cost of reverting back to the ancestral amino acid residues (*5, 9, 15, 30*). Therefore, we evaluated the difference between the fitness energies of the evolved descendant domain with (φ*) and without (φ) reversion to the ancestral residue: Δφ = φ* - φ (**Fig. 10b**). In this analysis, the descendant domains experienced 0.25 substitutions per site, which results in *p*-distances similar to the average maximum from ancestor to descendent observed within PF00001 ortho-domains. Δφ was computed for reversion at every site individually wherever the descendant sequence differed from the starting sequence.

**Figure 10.**
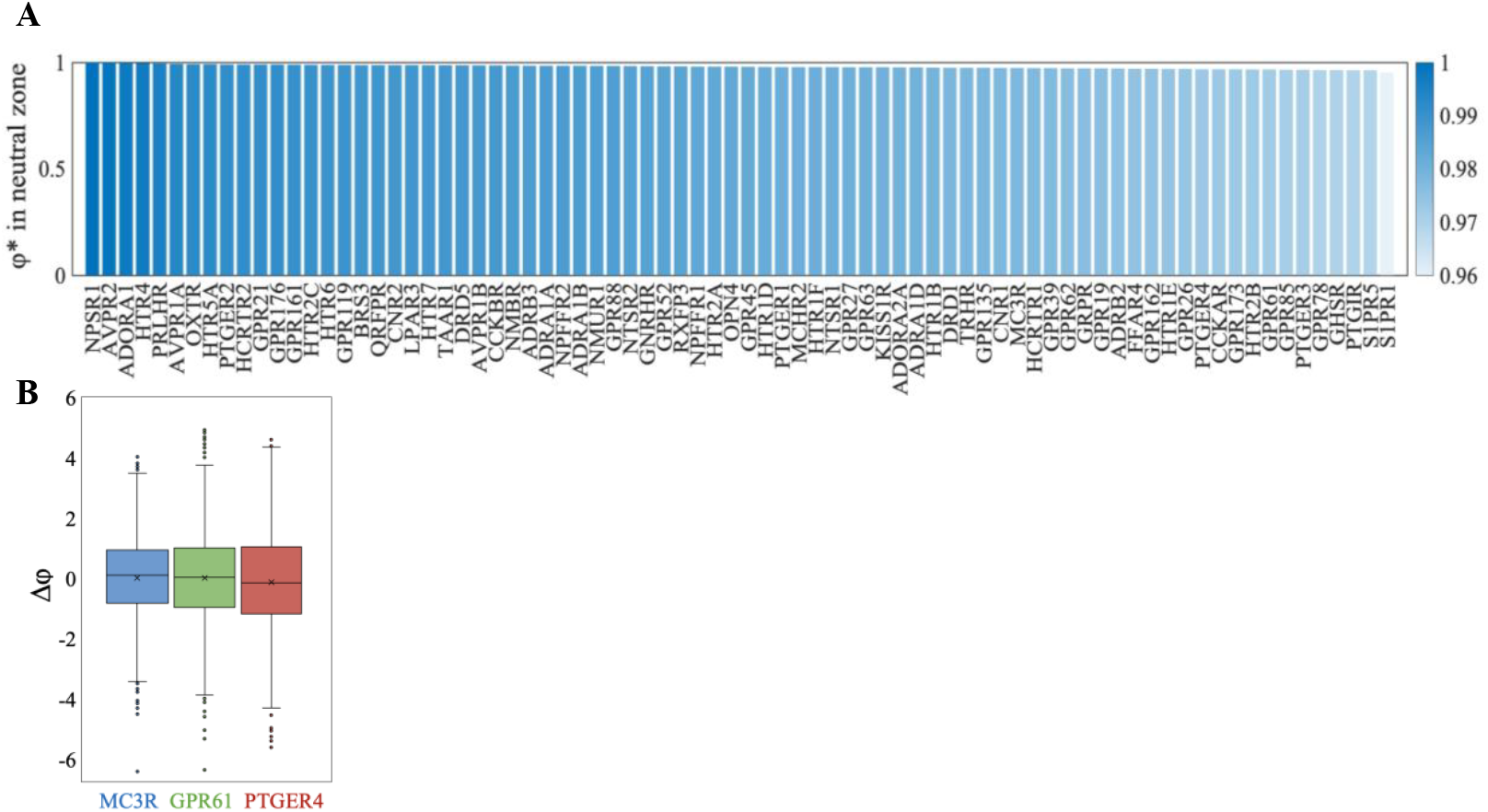
Energy cost (Δφ) of reverting single positions to ancestral residues. (**A**) Percentage of φ* values, descendent domains with single reversions to ancestral residues, which remained within the neutral zone for all PF00001 ortho-domains. On average, 97.92% of changes are neutral. (**B**) Change in energy (Δφ = φ* - φ) for three example ortho-domains (MC3R, GPR61, PTGER4) with low, median, and high energy (|φ|) respectively. Each plot is constructed for all positions which varied between ancestor and descendent across 10 replicates. All lineages were evolved to *d* = 0.25 substitutions per site.

For the PF00001 domain found in the GPR61 protein, evolving to *d* = 0.25 resulted in an average of 52 sequence differences between the ancestral and descendant domains. After single reversals, the average of Δφ was close to zero (0.04; *P* > 0.05) with nearly equal numbers of increases and decreases in φ* upon reversion to the ancestral amino acids. The average absolute deviation was also very small (|φ| = 1.36), a small fraction (0.24%) of the domain-specific φ for the PF00001 ortho-domain found in the GPR61 protein. In fact, only 3.36% of the φ* values fell outside the neutral zone, but they remained close (no more than 0.84% of total energy outside the zone). Therefore, telltale signs of entrenchment were not seen in neutral substitutions with epistasis for GPR61. Similar patterns were observed for two other domains of PF00001 found in the MC3R and PTGER4 proteins. The average of Δφ was again close to zero (0.03 - 0.11; **Fig. 10b**), and only a small percentage (2.65%, 3.25%) of φ* escaped the neutral zone. Similar trends in φ* were seen for all other domains in the PF00001 family (**Fig. 10a**); the ancestral residues generally remained energetically acceptable in the descendant domains. These results suggest a lack of significant entrenchment in neutral protein evolution with epistasis.

These entrenchment trends predict that neutral substitutions with epistasis do not significantly impact the substitution propensities of specific amino acids at individual sites, i.e., the ancestral and the descendant contexts have a limited impact on site-specific amino acid substitution probabilities. To assess amino acid propensities over time for a given position of an evolved domain (e.g., GPR61), we determined the amino acid with the highest propensity in the ancestral domain and then examined its propensity in the descendant domain (**Fig. 11a**). A linear relationship with a slope of 0.99 and a low dispersion (*R*^2^ = 0.99) shows that the most favorable amino-acid residue for substitution in the ancestral sequence (context) remained highly favored even in the descendant sequence (context) as many other sites contain neutral substitutions. Other amino-acid residues at each position also show similar patterns, as evident from sequence logos showing site-residue probabilities in the ancestral and descendent contexts (e.g., **Fig. 11b-c**). This pattern was confirmed in the analysis of other domains (**Fig. S5**).

**Figure 11.**
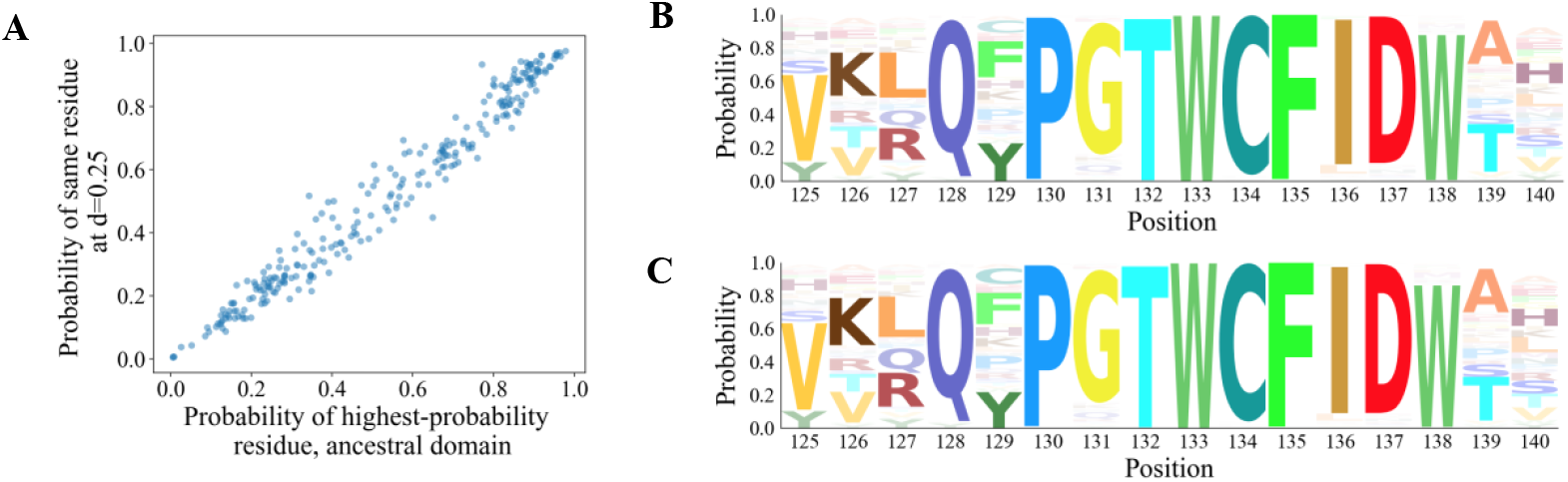
(**A**) For each site of the PF00001 domain (GPR61 protein; 268 amino acids), the highest-probability residue was found. Plotted are the initial probability of the residue, and the probability of the same residue after the domain has evolved to an evolutionary distance of *d* = 0.25 substitutions per site. A probability of 0 indicates an alignment gap in the domain. (**B–C**) Amino acid propensities for positions 125-140 of the domain of the ancestor (B) and the descendant (C) at *d* = 0.25 respectively. Site-residue probabilities were calculated for all amino acids using the PH model inferred from PF00001 homologs.

We also tested for the presence of entrenchment during neutral evolution with epistasis by examining Stokes Shift (*30*). Applying Stokes Shift analysis using PH energies (*5*), Δφ is computed by reversing an amino acid at a position of the descendent domain back to that found in the ancestral domain (Δφ_back_) and imposing an amino acid in the ancestral domain with that found in the same position of the descendent (Δφ_forward_). The sum of Δφ_forward_ and Δφ_back_ reflects the Stokes Shift during protein evolution, where the expectation is that Δφ_back_ + Δφ_forward_ = 0 when no Stokes shift occurs, i.e., Δφ_forward_ and Δφ_back_ have equal and opposite effects. Interestingly, in N×E evolution, the Stokes Shift was not significantly different from zero for every site at which the amino acid residue differed between the ancestral and descendant sequence (**Fig. 12**). This result confirms the lack of significant entrenchment. In addition, Stokes did not increase much even when we doubled the evolutionary distance (from 0.25 to 0.50), suggesting that even more distantly related domain orthologs may not suffer from significant entrenchment (**Fig. S6**).

**Figure 12.**
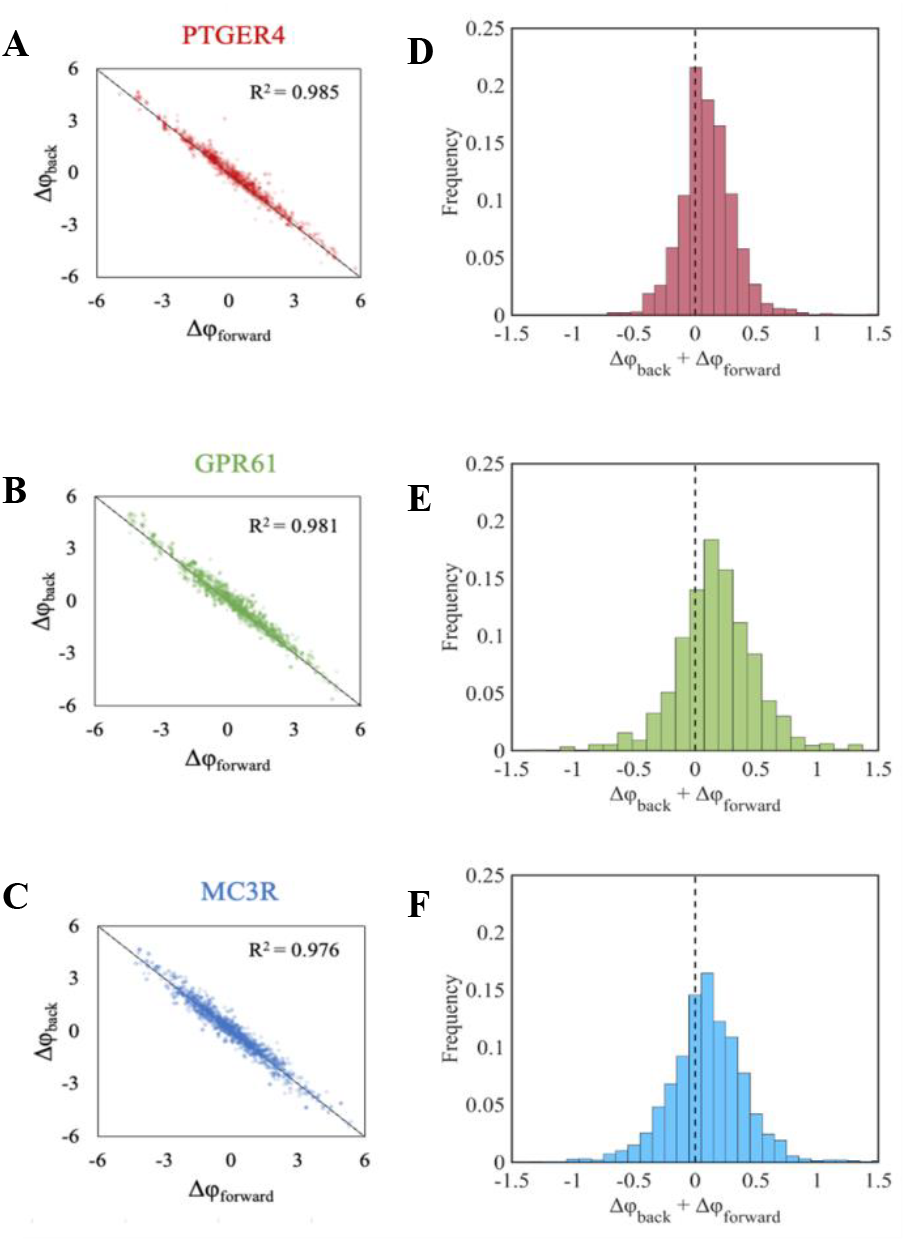
Stokes shift in NxE evolution. (**A–C**) Stokes shift plots of Δφ_forward_ and Δφ_back_ following the methods of de la Paz et al. (*5*). Each plot includes every site that differed between the ancestral and the descendent sequences (d = 0.25) across 30 different replicates. Calculations were performed on PF00001 ortho-domains with high |φ| (PTGER4), median φ (GPR61), and low |φ| (MC3R), respectively. (**D–F**) Histogram of Δφ_back_ + Δφ_forward_ for each site across all 30 replicates. The dotted line indicates Δφ_back_ + Δφ_forward_ = 0, i.e., no Stokes shift.

## Conclusions

We have introduced a neutral evolutionary framework in which the fitness of evolved domains is conserved, and epistasis dictates neutral substitutions. In this framework, purifying selection is imposed by the sequence context during the selection of an amino acid for substitution and by the constraint that its Hamiltonian Energy remains within the empirically-derived range for the ancestral domain. These requirements enable proper simulation of neutral protein evolution in which epistasis and functional conservation are considered.

Patterns of evolutionary rates among lineages show overdispersion of molecular clock, i.e., greater than expected variance of rates. These patterns of overdispersion are markedly different from those reported previously (*5, 31, 32*), as overdispersion plateaus rather than increasing continuously (**Fig. 8a**). Also, the highest overdispersion of the molecular clock is relatively modest in magnitude, e.g., 20% greater, on average, in simulated PF00001 ortho-domains. Although overdispersion persists over time, this persistence is not a consequence of the contingency of evolutionary rates, which would lead to consistently slower or faster rates in individual rates due to substitutions that occur early along the evolutionary trajectory. Instead, we find that the degree of overdispersion is a function of the proportion of evolutionary lineages in which evolutionary rates among sites have become significantly variable. As time progresses, site rates become variable in an increasing fraction of lineages, which causes increasing overdispersion of the molecular clock.

The most surprising pattern of neutral evolution with epistasis is the lack of entrenchment of amino-acid substitutions, which is a common side effect of epistasis. We found a lack of significant Stokes shift for amino-acid reversals, even at large timescales. Consequently, site-specific residue preferences remain largely conserved during neutral evolution with epistasis. Therefore, according to the N×E model, entrenchment is not an emergent property of epistasis in neutral protein evolution. Of course, models considering higher-order epistasis may lead to greater entrenchment (*33*). However, it is likely that PH models analyzed here will automatically account for some such scenarios (*34*). Other SEEC-like models, such as those using PH models fit to MSAs composed of highly varying within-population HIV sequence data, may be considered to model either non-neutral evolution or neutral evolution with a larger neutral range, and entrenchment is suggested to play a more important role (*8, 12*).

The patterns presented here suggest that a lack of contingency and absence of entrenchment may be considered null hypotheses under the neutral theory of molecular evolution with epistasis. With this null hypothesis, we are now developing novel approaches to sift through amino-acid substitutions and lineage-specific evolutionary rate changes for detecting bouts of non-neutral evolution in pursuit of adaptive patterns at individual sites, genes, and species.

## Acknowledgments Funding

National Science Foundation grant 1934848 (SK, VC, RL, AH, JT) National Institute of Health grant R35 GM139540-04 (SK) National Institute of Health grant R35 GM132090-04 (AH, RL)

## Author contributions

Conceptualization: SK, LS, VC, RL, AH

Methodology: LS, SK, RL

Investigation: LS, SC

Visualization: LS, SK

Funding acquisition: SK, VC, RL, AH, JT

Project administration: SK

Supervision: SK

Writing – original draft: LS

Writing – review & editing: SK, VC, RL, AH, JT

## Data and materials availability

Potts model parameters (couplings and local fields) required for NxE simulations were previously inferred by de la Paz et al. (*5*) and were made available at https://doi.org/10.5061/dryad.2ngf1vhj8. The SEEC simulation software is available at https://github.com/AlbertodelaPaz/SEEC. NxE simulations were performed by modifying the ‘*Probevolution*.*m’* software as provided in ‘*NxE*.*diff’*.

