## Supplementary Figures and Tables for "Entrenchment and contingency in neutral protein evolution with epistasis"

### Supplementary Materials

### Methods

*Constructing ortho-domain alignments*

We retrieved a total of 19 different protein domain family alignments analyzed in de la Paz et al. (*5*) and Figliuzzi et al. (*25*). For each family, we identified vertebrate orthologs by first searching for member sequences within the human proteome. We constructed a database of human proteins by selecting the human sequence from 19,077 UCSC 100-vertebrate alignments (*15*) and running individual BLAST searches for each of the domain families. Since we have a human to 100-vertebrate mapping within the alignments, it suffices to identify domains only within the human protein-coding sequences and then retrieve vertebrate orthologs of the same positions. We only consider hits with 100% sequence coverage and do not retain domains with insertions and deletions because the number of positions must match that of the family alignment in order to accurately calculate φ. Therefore, each ortho-domain alignment will contain up to 100 vertebrate species, but sequences may be removed when calculating the neutral range in energy (φ). After locating the positions of the domain in the vertebrate alignment, we removed all sites that were not part of the identified domain and sequences with missing data. We were able to construct over 300 ortho-domain alignments spanning a total of six different protein domain families (**Table S3**).

For domain families analyzed by the SEEC framework (*5*), the MSA and associated pairwise coupling (*e*) and local fields (*h*) matrices were extracted from the "Parameters_orig-x.mat" files made available by the authors, so no further PH model parameter inference was necessary (*https://doi.org/10.5061/dryad.2ngf1vhj8*). For the additional domain families analyzed in Figliuzzi et al. (*25*), we ran the bmDCA algorithm (*https://github.com/matteofigliuzzi/bmDCA*) on the provided MSAs of domain homologs to obtain *e* and *h* matrices.

*Potts Hamiltonian (PH) model construction*

Direct Coupling Analysis (DCA; *4*) is an umbrella term for the analysis of direct pairwise epistasis within a large MSA of homologous domains in order to produce a joint probability model, of which Potts Hamiltonian (PH) models are one variant. Some details of our simulations may depend on details of the PH models used in this analysis. Alternative PH inference algorithms make different approximations and yield somewhat different PH parameterizations when fit to the same MSA data. For instance, it was found that SEEC simulations differed when using PH models fit using mfDCA instead of bmDCA, and in some SEEC models a "selection temperature" can be tuned to adjust the simulation statistics (*24*). Many PH inference methods exist, but methods such as Mi3-GPU and bmDCA involving MCMC sampling of the model posterior distribution Boltzmann machine learning DCA (*25, 35, 36*) have been validated to capture the residue variability of the domain family with high precision. During inference of such PH models, clusters of orthologs with high sequence identity are downweighted, typically to have an effective count of 1 each. For PF00001 this left 6,593 effective sequences after weighting at an 80% sequence identity threshold, roughly representing an effective MSA of this many paralogous sequences.

The two components of the inferred PH model are a matrix of site-specific amino acid preferences (local fields; *h*) and a matrix of pairwise couplings (*e*) containing preferences for every pair of sites. From these components, Potts Hamiltonian Energy (PHE; φ) can be calculated for any sequence native to the protein domain family, as

$$P(\sigma_i = \alpha) = \frac{1}{Z} \exp \left\{ h_i(\alpha) + \sum_{j \neq i} e_{ij}(\alpha, \sigma_j) \right\} \quad (1)$$

In **Eq. 1**, *e* and *h* refer to the matrices as above, which are evaluated for each amino acid *σ* at site *i* ($\sigma_i$) and all possible pairs of sites ($\sigma_i$ , $\sigma_j$) for a sequence with *N* total positions.

*Simulating ortho-domain evolution*

To simulate ortho-domain evolution, we begin by following the procedure of the SEEC model (*5*). At each evolutionary step, we randomly sample a site along the protein by a uniform distribution where each site has an equal probability of being selected. We then sample an amino acid (including the probability of a synonymous change) from a conditional probability distribution (**Eq 2**), which describes the likelihood of finding each amino acid at the sampled position where the rest of the sequence is constant. This Boltzmann probability distribution treats each domain as a multivariate random variable and describes the probability of the sequence in relation to its family members, where *Z* is a normalizing constant.

$$\varphi = -\sum_{i=1}^{N-1} \sum_{j=i+1}^{N} e_{ij}(\sigma_i, \sigma_j) - \sum_{i=1}^{N} h_i(\sigma_i) \quad (2)$$

In other words, each possible residue (or an alignment gap character) is inserted into the sequence at the site being considered, and a distribution is constructed to find the instantaneous probabilities of substitution. While more probable sequences are favored, by the nature of the sampling procedure, there is still a chance of sampling less likely residues.

After a substitution is sampled, we calculate φ of the evolved domain. We assess if the substitution is predicted to be *neutral* as determined by the range in φ of sequences within the empirical ortho-domain alignment (see **Table S1** for PF00001 ortho-domain ranges). If φ falls outside the neutral range, we resample a site and residue by the above procedure until a permissible substitution is found. Although resampling occurs, when we track an evolutionary trajectory, only the permitted substitution and domain sequence are recorded.

*Justification for imposing a strict min-max boundary in φ (neutral zone)*

When we constructed alignments of ortho-domains in vertebrate species (UCSC 100-vertebrate alignments, see above), sequences had to be removed due to indels and missing data. Therefore, in many cases, the range in φ could not be calculated from all 100 sequences. Even when all data were available, the number of empirical sequences is limited to fully capture the statistical energies expected among domain orthologs. Therefore, for each of the 78 PF00001 ortho-domains, we calculated a normalized distribution of energies by finding the mean and variance of observations. We then found the percentage of the normalized distribution that was contained by the $\varphi_{min}$ and

$\varphi_{max}$ that were calculated from the empirical orthologs. On average, 92.1% of the distribution was contained (range 78.0–99.6%).

*Star phylogeny simulation procedure and calculating R*

We simulate star phylogenies following Fitch & Margoliash (*37*) and Kimura & Ohta (*26*) to avoid phylogenetic correlations among species. We begin with a sequence native to an ortho-domain alignment and simulate 100 lineages (replicates) emanating from the common ancestor. Lineages are evolved independently with all PH model parameters kept constant and the range in φ restricted to that of the empirical orthologs.

Star phylogenies have been classically used to test the molecular clock hypothesis by finding the number of substitutions that have accumulated in each species and computing *R* as the ratio of the variance to the mean. This calculation makes the underlying assumptions that the rate of all mutations and the selection intensity remain constant across species and over time. In N×E simulations, selection intensity remains constant because it is induced by the PH model and restriction in φ, both of which are constant among lineages in the tree. But, we must impose a constant mutation rate. If the mutation rate is constant among species, we expect that the *total number* of mutations in each lineage will be Poisson-distributed. Since each evolutionary step in N×E simulations either results in a synonymous or nonsynonymous substitution, it is sufficient to sample the number of evolutionary steps in each branch of the tree from a Poisson distribution with the same expectation. For each simulated tree, the mean number of evolutionary steps is chosen based on the purifying selection rate in the given ortho-domain and the desired evolutionary distance (e.g., $d$ = 0.25 or 0.5 substitutions per site).

We test these conditions in a control experiment without pairwise couplings (independent-site evolution) and a random evolution model in which all residues are equally likely to appear (uniform evolution). As expected, $R$=1 on average with minimal variance regardless of evolutionary distance. Therefore, any deviation from $R$=1 for N×E simulations will be due to pairwise epistasis.

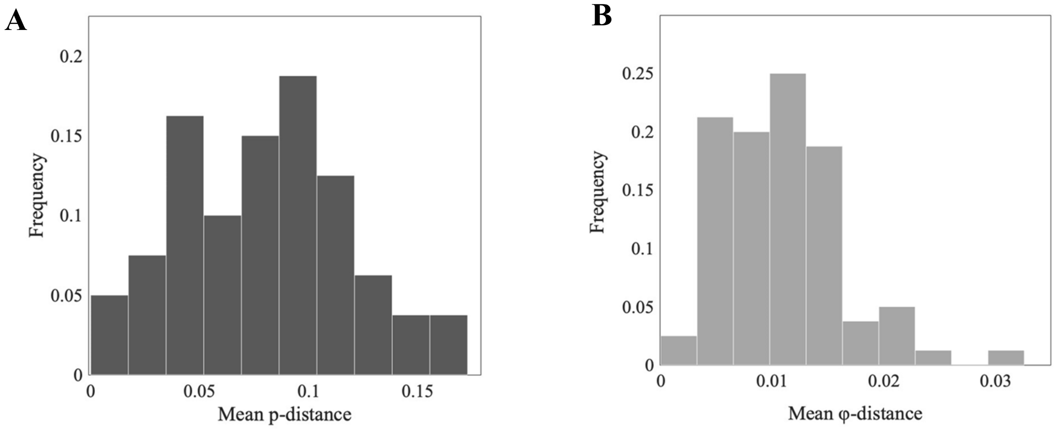


**Figure S1.** (**A**) Average *p*-distance for pairs of orthologs for each of the 78 variable PF00001 ortho-domain alignments (**B**) Average φ-distance for pairs of orthologs for each of the 78 variable PF00001 ortho-domain alignments

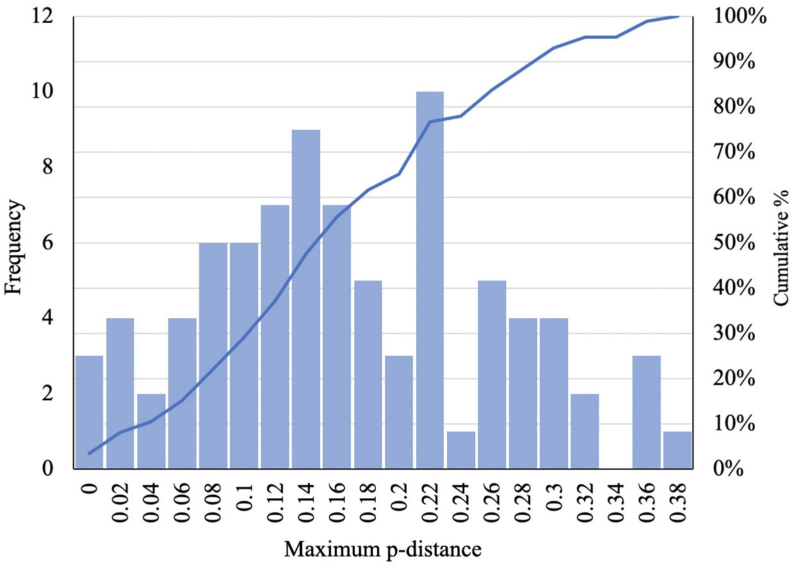


**Figure S2. Maximum *p*-distance within PF00001 ortho-domains.** For each of the 78 variable protein-coding genes with at least 1 copy of PF00001, we found the maximum *p*-distance among vertebrate orthologs. *p*-distance (Hamming distance) is calculated as the proportion of sites at which the two sequences are different in comparison to amino acid residues and alignment gaps.

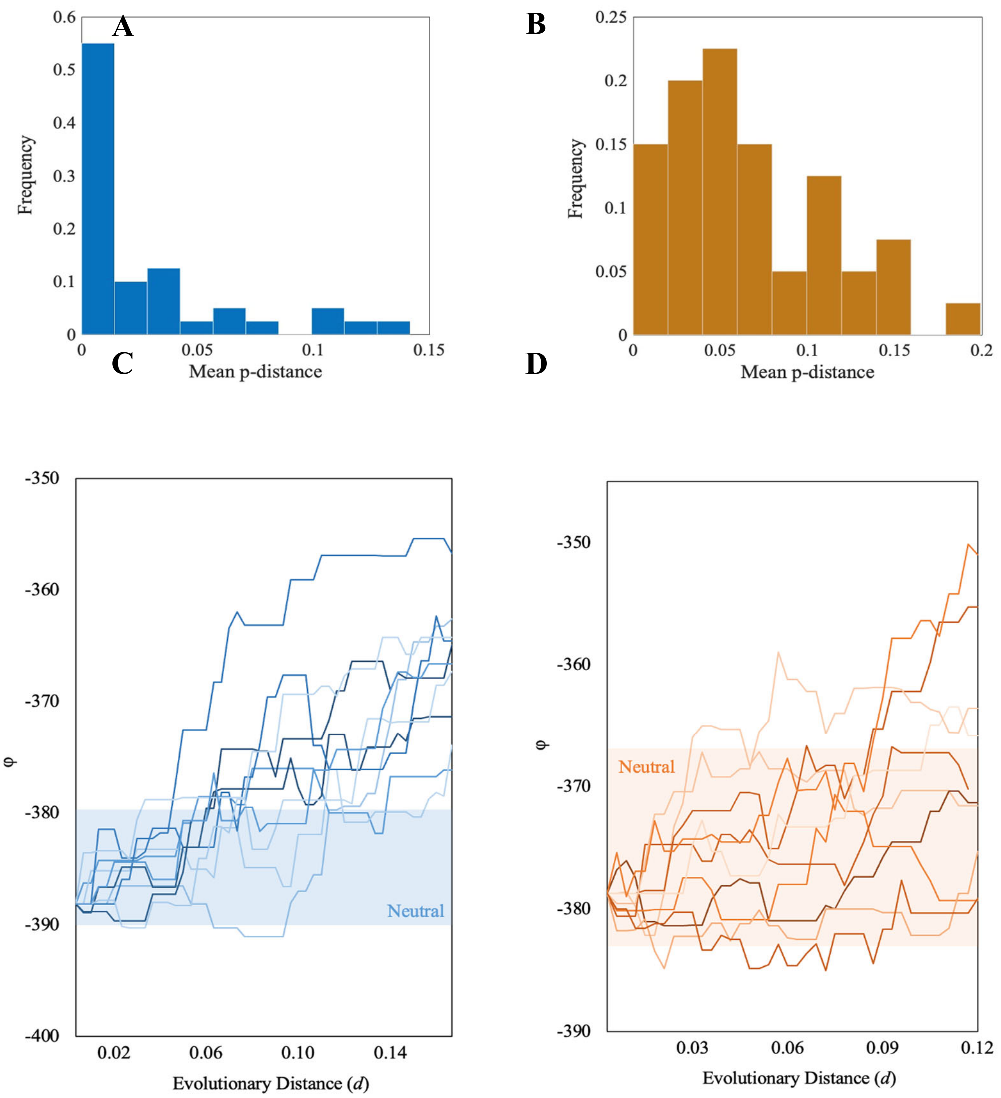


**Figure S3. Average *p*-distance among orthologs for PF0004 and PF00005 and φ trajectories.** (**A–B**) Average *p*-distance within ortho-domains (up to 100 vertebrate species) in each family (PF00004 and PF00005 respectively). (**C–D**) Trajectories of φ for single lineages evolved to *p*-distance = 0.1

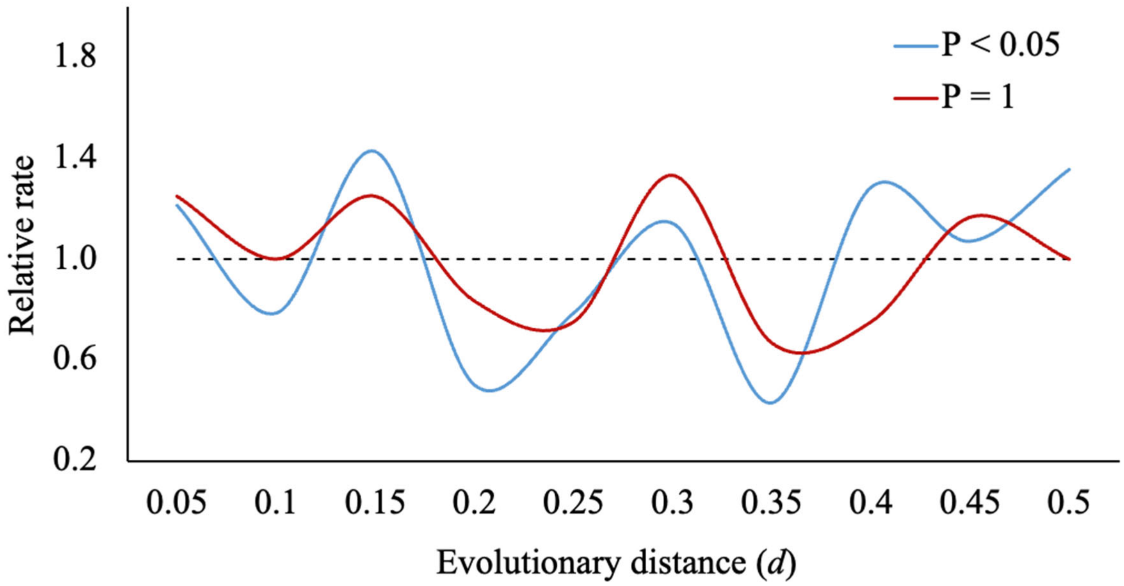


**Figure S4. Rate oscillation for a lineage that rejects the null hypothesis of no contingency.** Example of 2 lineages within the PTGER4 star tree: one that rejects the null hypothesis of no contingency ($P$ <0.05) and one that does not ($P$ = 1). Despite evolutionary rates being non-random, there is not a propensity for slow-evolving lineages to remain slow-evolving or vice versa. Instead, rates oscillate predictably from slow- to fast-evolving in the tree.

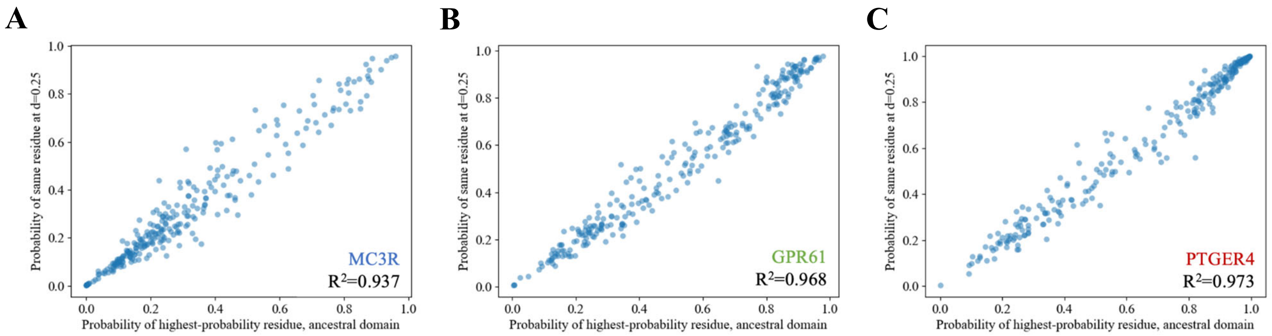


**Figure S5. Amino-acid propensities for ancestor and descendent at d = 0.25 (MC3R, GPR61 and PTGER4 proteins).** (**A**) Probability of highest-probability residue at ancestor versus probability of same residue at descendant evolved to $d$ = 0.25 substitutions per site (PF00001 domain found in the MC3R protein; $R^2$ = 0.937). (**B**) Probability of highest-probability residue at ancestor versus probability of same residue at descendant evolved to d = 0.25 substitutions per site (PF00001 domain found in the GPR61 protein; $R^2$ = 0.968). (**C**) Probability of highest-probability residue at ancestor versus probability of same residue at descendant evolved to d = 0.25 substitutions per site (PF00001 domain found in the PTGER4 protein; $R^2$ = 0.973).

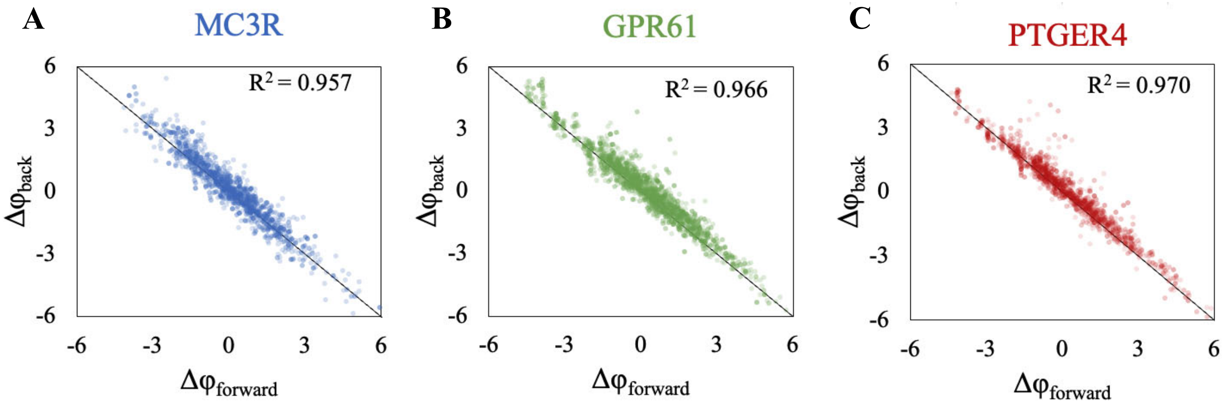


**Figure S6. Stokes shift at d = 0.5 substitutions per site.** (**A–C**) Stokes shift plots of $\Delta\varphi_{forward}$ and $\Delta\varphi_{back}$ following the methods of de la Paz et al. (*5*). Each plot includes every site which differed from ancestor to descendent (d = 0.5) across 30 different replicates. Calculations were done for PF00001 ortho-domains with low |φ| (MC3R), median φ (GPR61) and high |φ| (PTGER4), respectively.

%

**Table S1. Summary of statistical energies φ of PF00001 ortho-domains in 78 proteins.** Provided is the NMID of the alignment as retrieved from the UCSC 100-vertebrate alignments (*15*), gene name, and summary of φ values as calculated from the PH model provided by de la Paz et al. (*5*).

| Protein | NMID | Min φ | Max φ | Mean φ | φ Range |
|---|---|---|---|---|---|
| **PTGER4** | NM_000958 | -704.822158 | -693.732259 | -702.094 | 11.089899 |
| **TRHR** | NM_003301 | -720.473822 | -681.857128 | -692.069 | 38.616694 |
| **ADRB2** | NM_000024 | -688.696522 | -672.099206 | -683.130 | 16.597316 |
| **NMUR1** | NM_006056 | -692.52532 | -668.995698 | -680.908 | 23.529622 |
| **AVPR1A** | NM_000706 | -697.726517 | -648.722949 | -675.002 | 49.003568 |
| **ADORA2A** | NM_000675 | -691.759314 | -651.351909 | -674.511 | 40.407405 |
| **OXTR** | NM_000916 | -674.863732 | -655.212697 | -666.436 | 19.651035 |
| **ADRB1** | NM_000684 | -663.765905 | -663.765905 | -663.766 | 0 |
| **DRD5** | NM_000798 | -656.084801 | -648.21986 | -652.438 | 7.864941 |
| **AVPR1B** | NM_000707 | -664.938444 | -631.635449 | -650.842 | 33.302995 |
| **DRD1** | NM_000794 | -647.446488 | -630.19813 | -638.817 | 17.248358 |
| **TAAR1** | NM_138327 | -662.805856 | -604.864591 | -637.869 | 57.941265 |
| **OPN4** | NM_033282 | -640.418662 | -600.199931 | -631.237 | 40.218731 |
| **ADRA1B** | NM_000679 | -634.997964 | -619.338979 | -630.591 | 15.658985 |
| **NPFFR2** | NM_004885 | -640.013065 | -592.806112 | -628.540 | 47.206953 |
| **HTR2C** | NM_000868 | -633.636532 | -604.714545 | -628.298 | 28.921987 |
| **ADRA1A** | NM_000680 | -625.855902 | -601.952131 | -616.990 | 23.903771 |
| **HTR1F** | NM_001322209 | -624.6765 | -597.37581 | -615.701 | 27.30069 |
| **NPFFR1** | NM_022146 | -619.441135 | -602.015996 | -614.087 | 17.425139 |
| **ADRA1D** | NM_000678 | -616.486901 | -605.696747 | -613.438 | 10.790154 |
| **GPR26** | NM_153442 | -614.202656 | -598.716322 | -609.258 | 15.486334 |
| **HTR2A** | NM_000621 | -612.502925 | -588.297558 | -607.994 | 24.205367 |
| **ADRB3** | NM_000025 | -620.565076 | -589.878742 | -607.907 | 30.686334 |
| **HTR1D** | NM_000864 | -616.249939 | -590.971167 | -606.551 | 25.278772 |
| **HTR1B** | NM_000863 | -609.62804 | -594.598203 | -604.647 | 15.029837 |
| **AVPR2** | NM_000054 | -614.947365 | -585.17309 | -604.078 | 29.774275 |
| **HTR1E** | NM_000865 | -608.040499 | -597.286776 | -602.228 | 10.753723 |
| **ADORA1** | NM_000674 | -610.187239 | -557.330653 | -601.037 | 52.856586 |
| **GRPR** | NM_005314 | -601.819953 | -572.074781 | -592.932 | 29.745172 |
| **PTGIR** | NM_000960 | -584.612102 | -576.479235 | -581.304 | 8.132867 |
| **NTSR1** | NM_002531 | -594.282586 | -552.947237 | -576.998 | 41.335349 |
| **NMBR** | NM_002511 | -585.986498 | -558.498099 | -575.486 | 27.488399 |
| **GPR39** | NM_001508 | -585.256359 | -560.164315 | -574.270 | 25.092044 |
| **HTR2B** | NM_000867 | -597.06396 | -556.233717 | -572.160 | 40.830243 |
| **CCKAR** | NM_000730 | -581.542831 | -565.118093 | -571.481 | 16.424738 |
| **HTR7** | NM_019859 | -576.719138 | -541.186178 | -571.088 | 35.53296 |

| | | | | | |
|---|---|---|---|---|---|
| **GPR61** | NM_001393907 | -584.134718 | -562.129484 | -571.072 | 22.005234 |
| **CCKBR** | NM_176875 | -575.492677 | -562.978823 | -567.888 | 12.513854 |
| **HTR4** | NM_000870 | -564.726375 | -552.406872 | -556.948 | 12.319503 |
| **BRS3** | NM_001727 | -573.419086 | -536.063953 | -556.695 | 37.355133 |
| **CNR2** | NM_001841 | -569.277584 | -520.370321 | -555.216 | 48.907263 |
| **HCRTR2** | NM_001384272 | -550.753268 | -513.257146 | -543.166 | 37.496122 |
| **GNRHR** | NM_000406 | -553.39427 | -527.882972 | -539.351 | 25.511298 |
| **HTR5A** | NM_024012 | -546.59413 | -515.733056 | -535.287 | 30.861074 |
| **PTGER2** | NM_000956 | -554.553623 | -478.730826 | -535.117 | 75.822797 |
| **HTR6** | NM_000871 | -540.847969 | -521.088287 | -534.746 | 19.759682 |
| **KISS1R** | NM_032551 | -545.062762 | -524.354565 | -534.230 | 20.708197 |
| **S1PR1** | NM_001400 | -541.052965 | -516.56726 | -531.744 | 24.485705 |
| **PRLHR** | NM_004248 | -537.401726 | -496.769486 | -525.915 | 40.63224 |
| **HCRTR1** | NM_001525 | -530.763096 | -512.841604 | -525.529 | 17.921492 |
| **FFAR4** | NM_001195755 | -536.880945 | -495.35399 | -522.654 | 41.526955 |
| **CNR1** | NM_016083 | -521.955649 | -501.757257 | -517.554 | 20.198392 |
| **GPR78** | NM_080819 | -521.441843 | -507.128897 | -514.702 | 14.312946 |
| **S1PR5** | NM_030760 | -522.457616 | -499.238256 | -514.519 | 23.21936 |
| **GPR22** | NM_005295 | -511.392154 | -511.392154 | -511.392 | 0 |
| **GPR119** | NM_178471 | -522.826139 | -487.63306 | -509.890 | 35.193079 |
| **NPSR1** | NM_207172 | -515.718437 | -476.513518 | -505.977 | 39.204919 |
| **GPR161** | NM_001375883 | -509.407089 | -490.766993 | -503.160 | 18.640096 |
| **PTGER3** | NM_198719 | -509.799806 | -491.021836 | -501.179 | 18.77797 |
| **GPR45** | NM_007227 | -508.291431 | -470.964631 | -498.712 | 37.3268 |
| **QRFPR** | NM_198179 | -506.526335 | -479.68712 | -497.681 | 26.839215 |
| **GPR85** | NM_001146267 | -501.799348 | -485.280019 | -491.377 | 16.519329 |
| **PTGER1** | NM_000955 | -499.375806 | -475.335872 | -489.641 | 24.039934 |
| **GPR173** | NM_018969 | -488.977755 | -473.330441 | -486.032 | 15.647314 |
| **NTSR2** | NM_012344 | -493.140817 | -447.261111 | -480.036 | 45.879706 |
| **MCHR2** | NM_001040179 | -485.520459 | -455.722087 | -479.979 | 29.798372 |
| **GPR63** | NM_030784 | -485.100715 | -463.161765 | -478.738 | 21.93895 |
| **GPR21** | NM_005294 | -490.463064 | -467.361737 | -478.242 | 23.101327 |
| **GPR19** | NM_006143 | -482.315808 | -457.000409 | -474.319 | 25.315399 |
| **GHSR** | NM_198407 | -483.44077 | -459.034155 | -472.400 | 24.406615 |
| **GPR52** | NM_005684 | -474.629632 | -455.109244 | -468.295 | 19.520388 |
| **RXFP3** | NM_016568 | -471.716636 | -449.127921 | -463.732 | 22.588715 |
| **GPR27** | NM_018971 | -456.525267 | -448.917292 | -454.160 | 7.607975 |
| **GPR135** | NM_022571 | -453.779091 | -433.135369 | -444.387 | 20.643722 |
| **LPAR3** | NM_012152 | -432.815129 | -410.058791 | -425.050 | 22.756338 |
| **GPR88** | NM_022049 | -425.258179 | -415.368198 | -423.002 | 9.889981 |
| **GPR176** | NM_007223 | -426.513221 | -394.05316 | -420.666 | 32.460061 |

| | | | | | |
|---|---|---|---|---|---|
| **MC3R** | NM_019888 | -415.515959 | -400.307407 | -408.277 | 15.208552 |
| **GPR62** | NM_080865 | -408.897214 | -396.495607 | -404.366 | 12.401607 |
| **GPR162** | NM_019858 | -382.930692 | -378.225139 | -380.697 | 4.705553 |

**Table S2. Mean *R* values for star trees evolved to d = 0.5 substitutions per site, emanating from each of the 78 variable PF00001 ortho-domains.**

| Protein | Mean *R* |
|---|---|
| ADRB2 | 1.19950037 |
| ADRB3 | 1.07765429 |
| AVPR2 | 1.06494963 |
| GNRHR | 1.06714507 |
| HTR2A | 1.23772661 |
| ADORA1 | 1.0243695 |
| ADORA2A | 1.19075793 |
| ADRA1D | 1.27324121 |
| ADRA1B | 1.34544148 |
| ADRA1A | 1.29503672 |
| AVPR1A | 1.16103349 |
| AVPR1B | 1.10831582 |
| CCKAR | 1.21343401 |
| DRD1 | 1.40240792 |
| DRD5 | 1.45929035 |
| HTR1B | 1.11819071 |
| HTR1D | 1.18984456 |
| HTR1E | 1.18663543 |
| HTR2B | 1.26832556 |
| HTR2C | 1.21384758 |
| HTR4 | 1.03987682 |
| HTR6 | 1.10465527 |
| OXTR | 1.22072697 |
| PTGER1 | 1.11081073 |
| PTGER2 | 1.03056926 |
| PTGER4 | 1.2487132 |
| PTGIR | 1.17583453 |
| MCHR2 | 1.22142575 |

| | |
|---|---|
| GPR85 | 1.22961445 |
| FFAR4 | 1.14155954 |
| HTR1F | 1.07210193 |
| GPR161 | 1.01508891 |
| HCRTR2 | 1.12556558 |
| GPR61 | 1.02783205 |
| S1PR1 | 1.12874618 |
| GPR39 | 1.3249735 |
| HCRTR1 | 1.29962317 |
| BRS3 | 1.2460244 |
| CNR2 | 1.20032996 |
| NMBR | 1.12040204 |
| NTSR1 | 1.27203834 |
| TRHR | 1.31554914 |
| PRLHR | 1.11631746 |
| NPFFR2 | 1.13648622 |
| GPR21 | 1.32916204 |
| GRPR | 1.22726854 |
| GPR52 | 1.04253974 |
| NMUR1 | 1.19053517 |
| GPR19 | 1.08153177 |
| GPR176 | 1.11424318 |
| GPR45 | 1.18534026 |
| LPAR3 | 1.42515233 |
| NTSR2 | 1.21318888 |
| CNR1 | 1.27418078 |
| RXFP3 | 1.21823715 |
| GPR173 | 1.07472834 |
| GPR27 | 1.08249569 |
| GPR162 | 1.22547281 |
| HTR7 | 1.04220032 |

| | |
|---|---|
| MC3R | 1.08756273 |
| GPR88 | 1.17363904 |
| NPFFR1 | 1.17754722 |
| GPR135 | 1.33310312 |
| HTR5A | 1.16753649 |
| S1PR5 | 1.08499855 |
| GPR63 | 1.03335117 |
| KISS1R | 0.98193454 |
| OPN4 | 1.11271493 |
| GPR78 | 1.14242065 |
| GPR62 | 1.08749376 |
| TAAR1 | 1.26308816 |
| GPR26 | 1.23921249 |
| CCKBR | 1.20207787 |
| GPR119 | 1.1255193 |
| QRFPR | 1.03473346 |
| GHSR | 1.08414148 |
| PTGER3 | 1.25125275 |
| NPSR1 | 1.10083037 |

**Table S3. Summary of ortho-domains constructed for 6 different protein domain families, mean energy (φ), and maximum φ-distance among sequences.**

| Pfam ID | Pfam Short Name | Protein | NMID | Mean φ | Max φ-dist |
|---|---|---|---|---|---|
| PF00001 | 7tm_1 | ADRB2 | NM_000024 | -683.13 | 0.02 |
| PF00001 | 7tm_1 | ADRB3 | NM_000025 | -607.91 | 0.05 |
| PF00001 | 7tm_1 | AVPR2 | NM_000054 | -604.08 | 0.05 |
| PF00001 | 7tm_1 | GNRHR | NM_000406 | -539.35 | 0.05 |
| PF00001 | 7tm_1 | HTR2A | NM_000621 | -607.99 | 0.04 |
| PF00001 | 7tm_1 | ADORA1 | NM_000674 | -601.04 | 0.09 |
| PF00001 | 7tm_1 | ADORA2A | NM_000675 | -674.51 | 0.06 |
| PF00001 | 7tm_1 | ADRA1D | NM_000678 | -613.44 | 0.02 |
| PF00001 | 7tm_1 | ADRA1B | NM_000679 | -630.59 | 0.02 |
| PF00001 | 7tm_1 | ADRA1A | NM_000680 | -616.99 | 0.04 |
| PF00001 | 7tm_1 | ADRB1 | NM_000684 | -663.77 | 0 |
| PF00001 | 7tm_1 | AVPR1A | NM_000706 | -675 | 0.07 |
| PF00001 | 7tm_1 | AVPR1B | NM_000707 | -650.84 | 0.05 |
| PF00001 | 7tm_1 | CCKAR | NM_000730 | -571.48 | 0.03 |
| PF00001 | 7tm_1 | DRD1 | NM_000794 | -638.82 | 0.03 |
| PF00001 | 7tm_1 | DRD5 | NM_000798 | -652.44 | 0.01 |
| PF00001 | 7tm_1 | HTR1B | NM_000863 | -604.65 | 0.02 |
| PF00001 | 7tm_1 | HTR1D | NM_000864 | -606.55 | 0.04 |
| PF00001 | 7tm_1 | HTR1E | NM_000865 | -602.23 | 0.02 |
| PF00001 | 7tm_1 | HTR2B | NM_000867 | -572.16 | 0.07 |
| PF00001 | 7tm_1 | HTR2C | NM_000868 | -628.3 | 0.05 |
| PF00001 | 7tm_1 | HTR4 | NM_000870 | -556.95 | 0.02 |
| PF00001 | 7tm_1 | HTR6 | NM_000871 | -534.75 | 0.04 |
| PF00001 | 7tm_1 | OXTR | NM_000916 | -666.44 | 0.03 |
| PF00001 | 7tm_1 | PTGER1 | NM_000955 | -489.64 | 0.05 |
| PF00001 | 7tm_1 | PTGER2 | NM_000956 | -535.12 | 0.14 |
| PF00001 | 7tm_1 | PTGER4 | NM_000958 | -702.09 | 0.02 |
| PF00001 | 7tm_1 | PTGIR | NM_000960 | -581.3 | 0.01 |
| PF00001 | 7tm_1 | MCHR2 | NM_001040179 | -479.98 | 0.06 |
| PF00001 | 7tm_1 | GPR85 | NM_001146267 | -491.38 | 0.03 |
| PF00001 | 7tm_1 | FFAR4 | NM_001195755 | -522.65 | 0.08 |
| PF00001 | 7tm_1 | HTR1F | NM_001322209 | -615.7 | 0.04 |
| PF00001 | 7tm_1 | GPR161 | NM_001375883 | -503.16 | 0.04 |
| PF00001 | 7tm_1 | HCRTR2 | NM_001384272 | -543.17 | 0.07 |
| PF00001 | 7tm_1 | GPR61 | NM_001393907 | -571.07 | 0.04 |
| PF00001 | 7tm_1 | S1PR1 | NM_001400 | -531.74 | 0.05 |
| PF00001 | 7tm_1 | GPR39 | NM_001508 | -574.27 | 0.04 |
| PF00001 | 7tm_1 | HCRTR1 | NM_001525 | -525.53 | 0.03 |
| PF00001 | 7tm_1 | BRS3 | NM_001727 | -556.7 | 0.07 |
| PF00001 | 7tm_1 | CNR2 | NM_001841 | -555.22 | 0.09 |

| | | | | | |
|---|---|---|---|---|---|
| PF00001 | 7tm_1 | NMBR | NM_002511 | -575.49 | 0.05 |
| PF00001 | 7tm_1 | NTSR1 | NM_002531 | -577 | 0.07 |
| PF00001 | 7tm_1 | TRHR | NM_003301 | -692.07 | 0.06 |
| PF00001 | 7tm_1 | PRLHR | NM_004248 | -525.91 | 0.08 |
| PF00001 | 7tm_1 | NPFFR2 | NM_004885 | -628.54 | 0.08 |
| PF00001 | 7tm_1 | GPR21 | NM_005294 | -478.24 | 0.05 |
| PF00001 | 7tm_1 | GPR22 | NM_005295 | -511.39 | 0 |
| PF00001 | 7tm_1 | GRPR | NM_005314 | -592.93 | 0.05 |
| PF00001 | 7tm_1 | GPR52 | NM_005684 | -468.29 | 0.04 |
| PF00001 | 7tm_1 | NMUR1 | NM_006056 | -680.91 | 0.03 |
| PF00001 | 7tm_1 | GPR19 | NM_006143 | -474.32 | 0.05 |
| PF00001 | 7tm_1 | GPR176 | NM_007223 | -420.67 | 0.08 |
| PF00001 | 7tm_1 | GPR45 | NM_007227 | -498.71 | 0.07 |
| PF00001 | 7tm_1 | LPAR3 | NM_012152 | -425.05 | 0.05 |
| PF00001 | 7tm_1 | NTSR2 | NM_012344 | -480.04 | 0.1 |
| PF00001 | 7tm_1 | CNR1 | NM_016083 | -517.55 | 0.04 |
| PF00001 | 7tm_1 | RXFP3 | NM_016568 | -463.73 | 0.05 |
| PF00001 | 7tm_1 | GPR173 | NM_018969 | -486.03 | 0.03 |
| PF00001 | 7tm_1 | GPR27 | NM_018971 | -454.16 | 0.02 |
| PF00001 | 7tm_1 | GPR162 | NM_019858 | -380.7 | 0.01 |
| PF00001 | 7tm_1 | HTR7 | NM_019859 | -571.09 | 0.06 |
| PF00001 | 7tm_1 | MC3R | NM_019888 | -408.28 | 0.04 |
| PF00001 | 7tm_1 | GPR88 | NM_022049 | -423 | 0.02 |
| PF00001 | 7tm_1 | NPFFR1 | NM_022146 | -614.09 | 0.03 |
| PF00001 | 7tm_1 | GPR135 | NM_022571 | -444.39 | 0.05 |
| PF00001 | 7tm_1 | HTR5A | NM_024012 | -535.29 | 0.06 |
| PF00001 | 7tm_1 | S1PR5 | NM_030760 | -514.52 | 0.05 |
| PF00001 | 7tm_1 | GPR63 | NM_030784 | -478.74 | 0.05 |
| PF00001 | 7tm_1 | KISS1R | NM_032551 | -534.23 | 0.04 |
| PF00001 | 7tm_1 | OPN4 | NM_033282 | -631.24 | 0.06 |
| PF00001 | 7tm_1 | GPR78 | NM_080819 | -514.7 | 0.03 |
| PF00001 | 7tm_1 | GPR62 | NM_080865 | -404.37 | 0.03 |
| PF00001 | 7tm_1 | TAAR1 | NM_138327 | -637.87 | 0.09 |
| PF00001 | 7tm_1 | GPR26 | NM_153442 | -609.26 | 0.03 |
| PF00001 | 7tm_1 | CCKBR | NM_176875 | -567.89 | 0.02 |
| PF00001 | 7tm_1 | GPR119 | NM_178471 | -509.89 | 0.07 |
| PF00001 | 7tm_1 | QRFPR | NM_198179 | -497.68 | 0.05 |
| PF00001 | 7tm_1 | GHSR | NM_198407 | -472.4 | 0.05 |
| PF00001 | 7tm_1 | PTGER3 | NM_198719 | -501.18 | 0.04 |
| PF00001 | 7tm_1 | NPSR1 | NM_207172 | -505.98 | 0.08 |
| PF00004 | *AAA* | PEX6 | NM_000287 | -378.65 | 0.1 |
| PF00004 | *AAA* | PEX1 | NM_000466 | -371.69 | 0.06 |
| PF00004 | *AAA* | ATAD3C | NM_001039211 | -350.83 | 0.24 |

| | | | | | |
|---|---|---|---|---|---|
| PF00004 | *AAA* | BCS1L | NM_001079866 | -356.45 | 0.04 |
| PF00004 | *AAA* | ATAD3A | NM_001170535 | -353.29 | 0.07 |
| PF00004 | *AAA* | FIGNL1 | NM_001287492 | -386.54 | 0.05 |
| PF00004 | *AAA* | IQCA1L | NM_001304419 | -335.75 | 0.13 |
| PF00004 | *AAA* | FIGNL2 | NM_001384995 | -260.72 | 0.03 |
| PF00004 | *AAA* | KATNAL2 | NM_001387690 | -357.79 | 0.05 |
| PF00004 | *AAA* | NVL | NM_002533 | -367.42 | 0.04 |
| PF00004 | *AAA* | PSMC1 | NM_002802 | -363.68 | 0.02 |
| PF00004 | *AAA* | PSMC2 | NM_002803 | -340.02 | 0 |
| PF00004 | *AAA* | PSMC3 | NM_002804 | -205.18 | 0.01 |
| PF00004 | *AAA* | PSMC5 | NM_002805 | -372.87 | 0 |
| PF00004 | *AAA* | PSMC6 | NM_002806 | -324.6 | 0.02 |
| PF00004 | *AAA* | RFC1 | NM_002913 | -348.11 | 0.04 |
| PF00004 | *AAA* | RFC4 | NM_002916 | -368 | 0.04 |
| PF00004 | *AAA* | SPG7 | NM_003119 | -344.87 | 0.03 |
| PF00004 | *AAA* | ORC1 | NM_004153 | -342.59 | 0.05 |
| PF00004 | *AAA* | TRIP13 | NM_004237 | -391.08 | 0.03 |
| PF00004 | *AAA* | LONP1 | NM_004793 | -378.74 | 0.04 |
| PF00004 | *AAA* | VPS4B | NM_004869 | -154.63 | 0.07 |
| PF00004 | *AAA* | NSF | NM_006178 | -382.1 | 0.05 |
| PF00004 | *AAA* | PSMC4 | NM_006503 | -356.68 | 0 |
| PF00004 | *AAA* | AFG3L2 | NM_006796 | -373.38 | 0.03 |
| PF00004 | *AAA* | KATNA1 | NM_007044 | -375.08 | 0.03 |
| PF00004 | *AAA* | VCP | NM_007126 | -314.42 | 0.01 |
| PF00004 | *AAA* | RFC5 | NM_007370 | -189.29 | 0.03 |
| PF00004 | *AAA* | VPS4A | NM_013245 | -379.44 | 0.04 |
| PF00004 | *AAA* | ATAD2 | NM_014109 | -334 | 0.04 |
| PF00004 | *AAA* | YME1L1 | NM_014263 | -358.3 | 0.15 |
| PF00004 | *AAA* | SPAST | NM_014946 | -388.69 | 0.03 |
| PF00004 | *AAA* | ATAD2B | NM_017552 | -329.78 | 0.03 |
| PF00004 | *AAA* | FIGN | NM_018086 | -281.18 | 0.05 |
| PF00004 | *AAA* | WRNIP1 | NM_020135 | -374.96 | 0.07 |
| PF00004 | *AAA* | SPATA5L1 | NM_024063 | -320.95 | 0.06 |
| PF00004 | *AAA* | IQCA1 | NM_024726 | -184.92 | 0.12 |
| PF00004 | *AAA* | LONP2 | NM_031490 | -384.08 | 0.04 |
| PF00004 | *AAA* | ATAD3B | NM_031921 | -350.56 | 0.23 |
| PF00004 | *AAA* | KATNAL1 | NM_032116 | -376.25 | 0.03 |
| PF00004 | *AAA* | SPATA5 | NM_145207 | -359.87 | 0.05 |
| PF00004 | *AAA* | RFC2 | NM_181471 | -200.08 | 0.04 |
| PF00005 | ABC_tran | ABCD1 | NM_000033 | -376.49 | 0.08 |
| PF00005 | ABC_tran | ABCA4 | NM_000350 | -375.79 | 0.14 |
| PF00005 | ABC_tran | ABCC8 | NM_000352 | -373.05 | 0.06 |
| PF00005 | ABC_tran | ABCC2 | NM_000392 | -372.84 | 0.07 |

| | | | | | |
|---|---|---|---|---|---|
| PF00005 | ABC_tran | ABCB4 | NM_000443 | -370.59 | 0.05 |
| PF00005 | ABC_tran | CFTR | NM_000492 | -366.43 | 0.1 |
| PF00005 | ABC_tran | TAP1 | NM_000593 | -365.76 | 0.05 |
| PF00005 | ABC_tran | ABCF1 | NM_001025091 | -365.04 | 0.08 |
| PF00005 | ABC_tran | ABCA3 | NM_001089 | -359.8 | 0.14 |
| PF00005 | ABC_tran | ABCB5 | NM_001163941 | -356.37 | 0.09 |
| PF00005 | ABC_tran | ABCC6 | NM_001171 | -355.8 | 0.07 |
| PF00005 | ABC_tran | ABCC10 | NM_001198934 | -354.47 | 0.09 |
| PF00005 | ABC_tran | ABCB7 | NM_001271696 | -353.42 | 0.05 |
| PF00005 | ABC_tran | ABCA8 | NM_001288985 | -351.49 | 0.04 |
| PF00005 | ABC_tran | TAP2 | NM_001290043 | -349.37 | 0.07 |
| PF00005 | ABC_tran | ABCB1 | NM_001348946 | -339.56 | 0.06 |
| PF00005 | ABC_tran | ABCC11 | NM_001370497 | -339.23 | 0.09 |
| PF00005 | ABC_tran | ABCA10 | NM_001377321 | -337.89 | 0.04 |
| PF00005 | ABC_tran | ABCC12 | NM_001393797 | -337.65 | 0.03 |
| PF00005 | ABC_tran | ABCA2 | NM_001606 | -336.19 | 0.09 |
| PF00005 | ABC_tran | ABCD3 | NM_002858 | -333.86 | 0.08 |
| PF00005 | ABC_tran | ABCE1 | NM_002940 | -326.09 | 0.12 |
| PF00005 | ABC_tran | ABCB11 | NM_003742 | -325.67 | 0.06 |
| PF00005 | ABC_tran | ABCC3 | NM_003786 | -324.07 | 0.09 |
| PF00005 | ABC_tran | ABCG2 | NM_004827 | -321.85 | 0.1 |
| PF00005 | ABC_tran | ABCC1 | NM_004996 | -320.55 | 0.03 |
| PF00005 | ABC_tran | ABCD4 | NM_005050 | -318.54 | 0.08 |
| PF00005 | ABC_tran | ABCD2 | NM_005164 | -316.77 | 0.03 |
| PF00005 | ABC_tran | ABCA1 | NM_005502 | -305.5 | 0.03 |
| PF00005 | ABC_tran | ABCC5 | NM_005688 | -302.97 | 0.03 |
| PF00005 | ABC_tran | ABCB6 | NM_005689 | -300.44 | 0.06 |
| PF00005 | ABC_tran | ABCC4 | NM_005845 | -293.81 | 0.1 |
| PF00005 | ABC_tran | ABCB8 | NM_007188 | -293.61 | 0.08 |
| PF00005 | ABC_tran | ABCF2 | NM_007189 | -292.94 | 0.1 |
| PF00005 | ABC_tran | ABCB10 | NM_012089 | -292.84 | 0.13 |
| PF00005 | ABC_tran | ABCG1 | NM_016818 | -291.09 | 0.13 |
| PF00005 | ABC_tran | ABCF3 | NM_018358 | -289.37 | 0.11 |
| PF00005 | ABC_tran | ABCA7 | NM_019112 | -289.06 | 0.11 |
| PF00005 | ABC_tran | ABCB9 | NM_019625 | -285.59 | 0.07 |
| PF00005 | ABC_tran | ABCC9 | NM_020297 | -284.79 | 0.06 |
| PF00005 | ABC_tran | ABCG4 | NM_022169 | -275.56 | 0.05 |
| PF00005 | ABC_tran | ABCG5 | NM_022436 | -271.15 | 0.04 |
| PF00005 | ABC_tran | ABCG8 | NM_022437 | -241.23 | 0.05 |
| PF00005 | ABC_tran | ABCA9 | NM_080283 | -158.85 | 0.05 |
| PF00005 | ABC_tran | ABCA5 | NM_172232 | -77 | 0.16 |
| PF00005 | ABC_tran | ABCA12 | NM_173076 | -62.89 | 0.18 |
| PF00041 | fn3 | ITGB4 | NM_000213 | -151.66 | 0.09 |

| | | | | | |
|---|---|---|---|---|---|
| PF00041 | fn3 | MYBPC3 | NM_000256 | -159.31 | 0.11 |
| PF00041 | fn3 | TEK | NM_000459 | -126.49 | 0.09 |
| PF00041 | fn3 | NFASC | NM_001005388 | -170.46 | 0.12 |
| PF00041 | fn3 | MYBPHL | NM_001010985 | -165.14 | 0.16 |
| PF00041 | fn3 | NRCAM | NM_001037132 | -172.56 | 0.09 |
| PF00041 | fn3 | FNDC3A | NM_001079673 | -167.66 | 0.1 |
| PF00041 | fn3 | PTPRB | NM_001109754 | -131.6 | 0.14 |
| PF00041 | fn3 | SDK2 | NM_001144952 | -154.53 | 0.11 |
| PF00041 | fn3 | PTPRQ | NM_001145026 | -145.19 | 0.15 |
| PF00041 | fn3 | IGFN1 | NM_001164586 | -142.77 | 0.14 |
| PF00041 | fn3 | TTN | NM_001267550 | -190.48 | 0.1 |
| PF00041 | fn3 | DSCAM | NM_001389 | -147.99 | 0.13 |
| PF00041 | fn3 | NEO1 | NM_002499 | -145.11 | 0.08 |
| PF00041 | fn3 | PTPRD | NM_002839 | -174.45 | 0.17 |
| PF00041 | fn3 | PTPRF | NM_002840 | -173.09 | 0.13 |
| PF00041 | fn3 | PTPRS | NM_002850 | -172.94 | 0.05 |
| PF00041 | fn3 | ROBO1 | NM_002941 | -160.15 | 0.07 |
| PF00041 | fn3 | MYOM1 | NM_003803 | -165.32 | 0.18 |
| PF00041 | fn3 | MYOM2 | NM_003970 | -162.48 | 0.11 |
| PF00041 | fn3 | NCAM2 | NM_004540 | -71.64 | 0.2 |
| PF00041 | fn3 | IGDCC3 | NM_004884 | -141.27 | 0.09 |
| PF00041 | fn3 | CNTN2 | NM_005076 | -171.16 | 0.17 |
| PF00041 | fn3 | DCC | NM_005215 | -151.91 | 0.06 |
| PF00041 | fn3 | TIE1 | NM_005424 | -142.77 | 0.18 |
| PF00041 | fn3 | CHL1 | NM_006614 | -169.24 | 0.22 |
| PF00041 | fn3 | CNTN5 | NM_014361 | -158.45 | 0.08 |
| PF00041 | fn3 | EPHB2 | NM_017449 | -168.64 | 0.14 |
| PF00041 | fn3 | IGDCC4 | NM_020962 | -159.35 | 0.15 |
| PF00041 | fn3 | FNDC3B | NM_022763 | -168.46 | 0.08 |
| PF00041 | fn3 | FNDC1 | NM_032532 | -137.39 | 0.12 |
| PF00041 | fn3 | CNTFR | NM_147164 | -130.18 | 0.08 |
| PF00041 | fn3 | FNDC5 | NM_153756 | -133.65 | 0.06 |
| PF00041 | fn3 | PRTG | NM_173814 | -140.57 | 0.08 |
| PF00153 | Mito_carr | SLC25A20 | NM_000387 | -208.62 | 0.07 |
| PF00153 | Mito_carr | SLC25A30 | NM_001010875 | -200.5 | 0.03 |
| PF00153 | Mito_carr | SLC25A29 | NM_001039355 | -197.36 | 0.02 |
| PF00153 | Mito_carr | SLC25A36 | NM_001104647 | -195.79 | 0.05 |
| PF00153 | Mito_carr | SLC25A19 | NM_001126121 | -193.8 | 0.15 |
| PF00153 | Mito_carr | SLC25A4 | NM_001151 | -191.48 | 0.07 |
| PF00153 | Mito_carr | SLC25A5 | NM_001152 | -190.04 | 0.04 |
| PF00153 | Mito_carr | SLC25A22 | NM_001191061 | -188.64 | 0.1 |
| PF00153 | Mito_carr | SLC25A14 | NM_001282195 | -186.79 | 0.06 |
| PF00153 | Mito_carr | SLC25A6 | NM_001636 | -128.79 | 0.06 |

| | | | | | |
|---|---|---|---|---|---|
| PF00153 | Mito_carr | UCP3 | NM_003356 | -117.26 | 0.07 |
| PF00153 | Mito_carr | SLC25A11 | NM_003562 | -115.84 | 0.08 |
| PF00153 | Mito_carr | SLC25A27 | NM_004277 | -113.07 | 0.14 |
| PF00153 | Mito_carr | SLC25A1 | NM_005984 | -112.5 | 0.1 |
| PF00153 | Mito_carr | SLC25A10 | NM_012140 | -109.96 | 0.08 |
| PF00153 | Mito_carr | SLC25A24 | NM_013386 | -80.41 | 0.25 |
| PF00153 | Mito_carr | SLC25A15 | NM_014252 | -74.29 | 0.14 |
| PF00153 | Mito_carr | SLC25A37 | NM_016612 | -73.61 | 0.21 |
| PF00153 | Mito_carr | SLC25A38 | NM_017875 | -64.12 | 0.17 |
| PF00153 | Mito_carr | UCP1 | NM_021833 | -64.12 | 0.22 |
| PF00153 | Mito_carr | SLC25A23 | NM_024103 | -58.03 | 0.12 |
| PF00153 | Mito_carr | SLC25A21 | NM_030631 | -57.96 | 0.16 |
| PF00153 | Mito_carr | SLC25A32 | NM_030780 | -57.76 | 0.08 |
| PF00153 | Mito_carr | SLC25A28 | NM_031212 | -54.49 | 0.3 |
| PF00153 | Mito_carr | SLC25A31 | NM_031291 | -52.02 | 0.3 |
| PF00153 | Mito_carr | SLC25A18 | NM_031481 | -47.66 | 0.19 |
| PF00153 | Mito_carr | SLC25A33 | NM_032315 | -46.07 | 0.41 |
| PF00153 | Mito_carr | SLC25A16 | NM_152707 | -44.21 | 0.28 |
| PF00153 | Mito_carr | SLC25A42 | NM_178526 | -40.58 | 0 |
| PF00271 | Helicase_C | BLM | NM_000057 | -299.1 | 0.04 |
| PF00271 | Helicase_C | ERCC6 | NM_000124 | -277.54 | 0.06 |
| PF00271 | Helicase_C | WRN | NM_000553 | -268.76 | 0.05 |
| PF00271 | Helicase_C | CHD3 | NM_001005273 | -299.63 | 0.06 |
| PF00271 | Helicase_C | DDX59 | NM_001031725 | -228.5 | 0.07 |
| PF00271 | Helicase_C | SHPRH | NM_001042683 | -229.44 | 0.06 |
| PF00271 | Helicase_C | CHD8 | NM_001170629 | -301.35 | 0.03 |
| PF00271 | Helicase_C | CHD1 | NM_001270 | -300.98 | 0.03 |
| PF00271 | Helicase_C | CHD2 | NM_001271 | -299.61 | 0.04 |
| PF00271 | Helicase_C | CHD4 | NM_001273 | -300.57 | 0.03 |
| PF00271 | Helicase_C | SMARCA1 | NM_001282874 | -305.28 | 0.05 |
| PF00271 | Helicase_C | DDX46 | NM_001300860 | -253.67 | 0.04 |
| PF00271 | Helicase_C | CHD9 | NM_001308319 | -296.04 | 0.05 |
| PF00271 | Helicase_C | DDX3X | NM_001356 | -300.07 | 0.02 |
| PF00271 | Helicase_C | EIF4A1 | NM_001416 | -298.87 | 0.06 |
| PF00271 | Helicase_C | EIF4A2 | NM_001967 | -303.25 | 0.03 |
| PF00271 | Helicase_C | RECQL | NM_002907 | -294.28 | 0.05 |
| PF00271 | Helicase_C | SMARCA2 | NM_003070 | -293.15 | 0.03 |
| PF00271 | Helicase_C | HLTF | NM_003071 | -252.33 | 0.06 |
| PF00271 | Helicase_C | SMARCA4 | NM_003072 | -297.34 | 0.02 |
| PF00271 | Helicase_C | SUPV3L1 | NM_003171 | -235.03 | 0.04 |
| PF00271 | Helicase_C | RAD54L | NM_003579 | -276.68 | 0.04 |
| PF00271 | Helicase_C | TTF2 | NM_003594 | -247.12 | 0.07 |
| PF00271 | Helicase_C | SMARCA5 | NM_003601 | -236.77 | 0.05 |

| | | | | | |
|---|---|---|---|---|---|
| PF00271 | Helicase_C | BTAF1 | NM_003972 | -270.76 | 0.02 |
| PF00271 | Helicase_C | RECQL5 | NM_004259 | -280.16 | 0.06 |
| PF00271 | Helicase_C | RECQL4 | NM_004260 | -106.06 | 0.14 |
| PF00271 | Helicase_C | CHD1L | NM_004284 | -268.4 | 0.05 |
| PF00271 | Helicase_C | DDX5 | NM_004396 | -88.56 | 0.06 |
| PF00271 | Helicase_C | DDX6 | NM_004397 | -66.89 | 0.07 |
| PF00271 | Helicase_C | DDX10 | NM_004398 | -223.36 | 0.08 |
| PF00271 | Helicase_C | DDX39B | NM_004640 | -78.61 | 0.07 |
| PF00271 | Helicase_C | DDX3Y | NM_004660 | -299.93 | 0.04 |
| PF00271 | Helicase_C | DDX21 | NM_004728 | -279.1 | 0.1 |
| PF00271 | Helicase_C | DDX23 | NM_004818 | -263.73 | 0.05 |
| PF00271 | Helicase_C | DDX1 | NM_004939 | -247.1 | 0.04 |
| PF00271 | Helicase_C | DDX39A | NM_005804 | -78.07 | 0.06 |
| PF00271 | Helicase_C | DDX17 | NM_006386 | -91.14 | 0.04 |
| PF00271 | Helicase_C | SRCAP | NM_006662 | -278.39 | 0.35 |
| PF00271 | Helicase_C | DDX18 | NM_006773 | -81.54 | 0.07 |
| PF00271 | Helicase_C | ASCC3 | NM_006828 | -262.39 | 0.1 |
| PF00271 | Helicase_C | DDX52 | NM_007010 | -99.4 | 0.14 |
| PF00271 | Helicase_C | DDX20 | NM_007204 | -39.83 | 0.37 |
| PF00271 | Helicase_C | DDX19B | NM_007242 | -266.24 | 0.05 |
| PF00271 | Helicase_C | RAD54B | NM_012415 | -276.32 | 0.07 |
| PF00271 | Helicase_C | DDX25 | NM_013264 | -261.78 | 0.16 |
| PF00271 | Helicase_C | SMARCAL1 | NM_014140 | -249.88 | 0.09 |
| PF00271 | Helicase_C | DDX58 | NM_014314 | -209.3 | 0.11 |
| PF00271 | Helicase_C | DHX34 | NM_014681 | -287.87 | 0.06 |
| PF00271 | Helicase_C | EIF4A3 | NM_014740 | -280.11 | 0.02 |
| PF00271 | Helicase_C | RAD54L2 | NM_015106 | -255.93 | 0.02 |
| PF00271 | Helicase_C | EP400 | NM_015409 | -53.93 | 0.27 |
| PF00271 | Helicase_C | CHD5 | NM_015557 | -297.71 | 0.06 |
| PF00271 | Helicase_C | DDX41 | NM_016222 | -210.75 | 0.04 |
| PF00271 | Helicase_C | DDX47 | NM_016355 | -279.38 | 0.05 |
| PF00271 | Helicase_C | INO80 | NM_017553 | -279.83 | 0.01 |
| PF00271 | Helicase_C | ERCC6L | NM_017669 | -254.26 | 0.08 |
| PF00271 | Helicase_C | CHD7 | NM_017780 | -302.78 | 0.12 |
| PF00271 | Helicase_C | DDX27 | NM_017895 | -263.97 | 0.11 |
| PF00271 | Helicase_C | HELLS | NM_018063 | -289.77 | 0.05 |
| PF00271 | Helicase_C | DDX19A | NM_018332 | -266.36 | 0.04 |
| PF00271 | Helicase_C | DDX28 | NM_018380 | -212.42 | 0.04 |
| PF00271 | Helicase_C | DDX43 | NM_018665 | -244.15 | 0.06 |
| PF00271 | Helicase_C | DDX49 | NM_019070 | -244.57 | 0.05 |
| PF00271 | Helicase_C | SMARCAD1 | NM_020159 | -254.15 | 0.05 |
| PF00271 | Helicase_C | ERCC6L2 | NM_020207 | -244.94 | 0.06 |
| PF00271 | Helicase_C | DDX24 | NM_020414 | -72.64 | 0.12 |

| | | | | | |
|---|---|---|---|---|---|
| PF00271 | Helicase_C | DDX55 | NM_020936 | -213.06 | 0.07 |
| PF00271 | Helicase_C | FANCM | NM_020937 | -269.92 | 0.06 |
| PF00271 | Helicase_C | IFIH1 | NM_022168 | -232.61 | 0.05 |
| PF00271 | Helicase_C | DDX50 | NM_024045 | -271.78 | 0.04 |
| PF00271 | Helicase_C | DDX54 | NM_024072 | -248.89 | 0.07 |
| PF00271 | Helicase_C | DDX4 | NM_024415 | -274.02 | 0.07 |
| PF00271 | Helicase_C | ZRANB3 | NM_032143 | -254.67 | 0.09 |
| PF00271 | Helicase_C | CHD6 | NM_032221 | -299.01 | 0.03 |
| PF00271 | Helicase_C | DDX51 | NM_175066 | -228.6 | 0.07 |
| PF00271 | Helicase_C | DICER1 | NM_177438 | -198.51 | 0.02 |
| PF00271 | Helicase_C | DDX53 | NM_182699 | -178.59 | 0.26 |
| PF00271 | Helicase_C | DDX42 | NM_203499 | -262.43 | 0.02 |
